# De novo design of phosphorylation-induced protein switches for synthetic signaling in cells

**DOI:** 10.1101/2025.09.10.675034

**Authors:** Stephen Buckley, Yangyang Miao, Mubarak Idris, Pao-Wan Lee, Leo Scheller, Roland Riek, Sebastian J. Maerkl, Luciano A. Abriata, Bruno E. Correia

**Author notes:** These authors contributed equally (Stephen Buckley, Yangyang Miao).

## Abstract

A hallmark of living systems is their ability to respond and adapt to many types of exogenous and endogenous cues. At the core of this capability lie the evolved mechanisms of cellular signaling which perform the relay of biochemical signals throughout the cell. Much of this core signaling functionality in living systems is afforded by the ability of proteins to change their chemical and structural “status”, by populating distinct states in conformational landscapes, undergoing post-translational modifications that alter these populations, and dynamically interacting with other cellular partners. Despite the remarkable advances we have witnessed in computational protein design, it remains an outstanding challenge to rationally design many of these aspects that are at the core of biological function. Here, we set out to design a minimal signaling network triggered by phosphorylation, which relies on de novo components that sample low-populated conformational states to control a designed protein interaction. In these designed components we implemented principles of signal amplification, conformational heterogeneity and molecular recognition which are ubiquitously used by nature to sustain biological function in cells. The designed proteins were biochemically and structurally characterized and ultimately are functional in cell-based systems where they regulate the transcription of reporter proteins in a phosphorylation-dependent manner. Overall, this work lays the foundation for designing synthetic signaling cascades using de novo protein components, which could be important to enhance our understanding of natural systems and for relevant applications in synthetic biology.

## Introduction

Cells, as the fundamental units in biology, have evolved intricate signaling pathways to sense extra and intracellular information and transmit biochemical signals through their milieu. Generally, distinct inputs result in changes in transcriptional programs to respond to the cell’s environment. To relay biochemical signals, many natural proteins are intrinsically dynamic, sampling both major and minor states within their conformational landscape. This dynamic behavior enables inducible mechanisms, such as allostery or post-translational modifications (PTMs), to regulate their conformation and function^1^. Such conformational changes often trigger downstream events, as seen in signaling enzymes, including kinases. For example, the inactive state of some receptor tyrosine kinases (RTKs), such as fibroblast growth factor receptors, samples a minor active conformation in addition to the more populated inactive conformation at equilibrium^2^. Upon extracellular ligand binding, two basally active RTKs are brought into proximity, facilitating trans-phosphorylation and shifting their conformational landscapes to predominantly populate the active state. This mechanism represents one way by which cells link an extracellular input to an intracellular signal. Additionally, phosphorylation can modulate conformational populations to regulate protein-protein interactions (PPIs), thereby controlling signal propagation. The cAMP response element-binding protein (CREB) undergoes a conformational change upon phosphorylation by protein kinase A (PKA), stabilizing a conformation that facilitates binding to CREB binding protein (CBP), resulting in a phosphorylation-dependent PPI^3^. These interactions can influence the target protein’s activity, localization, stability, or facilitate their interaction with another protein^4,5^. The ability to precisely regulate intracellular signals is vital to a cell’s survival, as the dysregulation of these signals can lead to disease states such as cancer^6,7^. Such signaling mechanisms are present across all kingdoms of life, where, in general, more complex organisms require more sophisticated signaling pathways to support their function. In simple organisms such as prokaryotes, the most typical example is the two-component system, composed of a sensor kinase and response regulator. These phosphorelay systems are thought to have ultimately evolved into the multi-step kinase cascades found in eukaryotes^8,9^.

Recently, great strides have been made in the field of protein engineering, such as algorithms that can accurately predict a structure based on a sequence^10,11^ or a sequence based on a structure^12^. However, to date, de novo designed proteins have largely been limited to the design of single-state, stable protein folds, with few examples existing for designed proteins with built-in dynamics due to an input stimulus^13,14,15,16^. Designing de novo proteins that react to stimuli provides an opportunity to mimic the dynamic nature of natural proteins and therefore replicate their inducible functions.

An enduring goal for synthetic biology is to design novel regulatory circuits within cells, conferring new functions. One mechanism used by proteins to regulate signals in such circuits is autoinhibition, where an inhibitory region of a protein blocks a functional site. For example, the Dbl Homology domain (a guanine nucleotide exchange factor for Rho GTPases) of the Vav1 protein exists in equilibrium between a major inhibited state, where an N-terminal helix blocks its catalytic site, and a minor activated state, where the N-terminal helix is displaced^17,18^. This minor state transiently exposes a previously buried phosphorylation site, granting accessibility to a kinase. Once phosphorylated, the equilibrium shifts to favor the active state, thereby regulating Vav1’s ability to propagate signals in a phosphorylation-dependent manner. Recently, synthetic cellular signaling systems have incorporated phosphorylation as a mechanism to regulate pathways, and therefore their response to a specific input^19,20^. While some progress has been made in designing phosphorylation-induced changes in de novo proteins^13^, to our knowledge, de novo protein components have not yet been implemented to interact in a phosphorylation-dependent manner. Such a system would mimic natural signaling proteins, where this phosphorylation-inducible interaction could propagate a biochemical signal in a cell. The ability to design de novo proteins capable of transmitting a biochemical signal would open the possibility for building more complex modular synthetic signaling cascades, allowing for the development of functionally-relevant cellular logic gates.

To explore protein engineering’s potential application to designing such systems, we set out to incorporate phosphorylation-sensitive sites into de novo proteins, switching their energetically favorable conformation in a phosphorylation-induced manner. We aimed to utilize these phosphorylation-induced protein switches to serve as synthetic signaling proteins in artificial cellular signaling cascades. To accomplish this, de novo folds were selected and engineered to incorporate a phospho-switch motif containing a tobacco etch virus (TEV) protease cleavage site and a protein kinase A (PKA) phosphorylation consensus sequence (RRxS), both of which are inaccessible in the non-phosphorylated “closed” state (Figure 1A). Once phosphorylated, the bulky dianionic phosphate group switches the energetically favorable state to an “open” conformation, exposing the TEV cleavage site. Through this design strategy we aimed to investigate which sequences encode the conformational dynamics necessary to support effective phosphorylation-responsive switching. Such switching is dependent on stabilizing the non-phosphorylated state in the closed conformation, thereby minimizing the population of the open conformation. However, some intrinsic dynamics must be retained to allow the kinase to access the shielded phosphorylation motif. To characterize the conformational landscape of the phospho-switches we performed detailed NMR studies. We demonstrated the generalizability of this approach by applying it to four de novo folds, including a novel fold. Furthermore, to relay the biochemical signal, we designed and screened binders specific to the open or phosphorylated state for one of the protein switches. Validated binders were integrated into cellular signaling systems, where the phosphorylation-induced PPI regulated the activity of a split transcription factor, driving expression of a reporter protein. By designing a simple phosphorylation-dependent PPI based on de novo protein components, we make a step towards designing more complex synthetic cellular signaling pathways with potential applications in cellular logic gates for therapeutics and synthetic biology tools.

**Figure 1:**
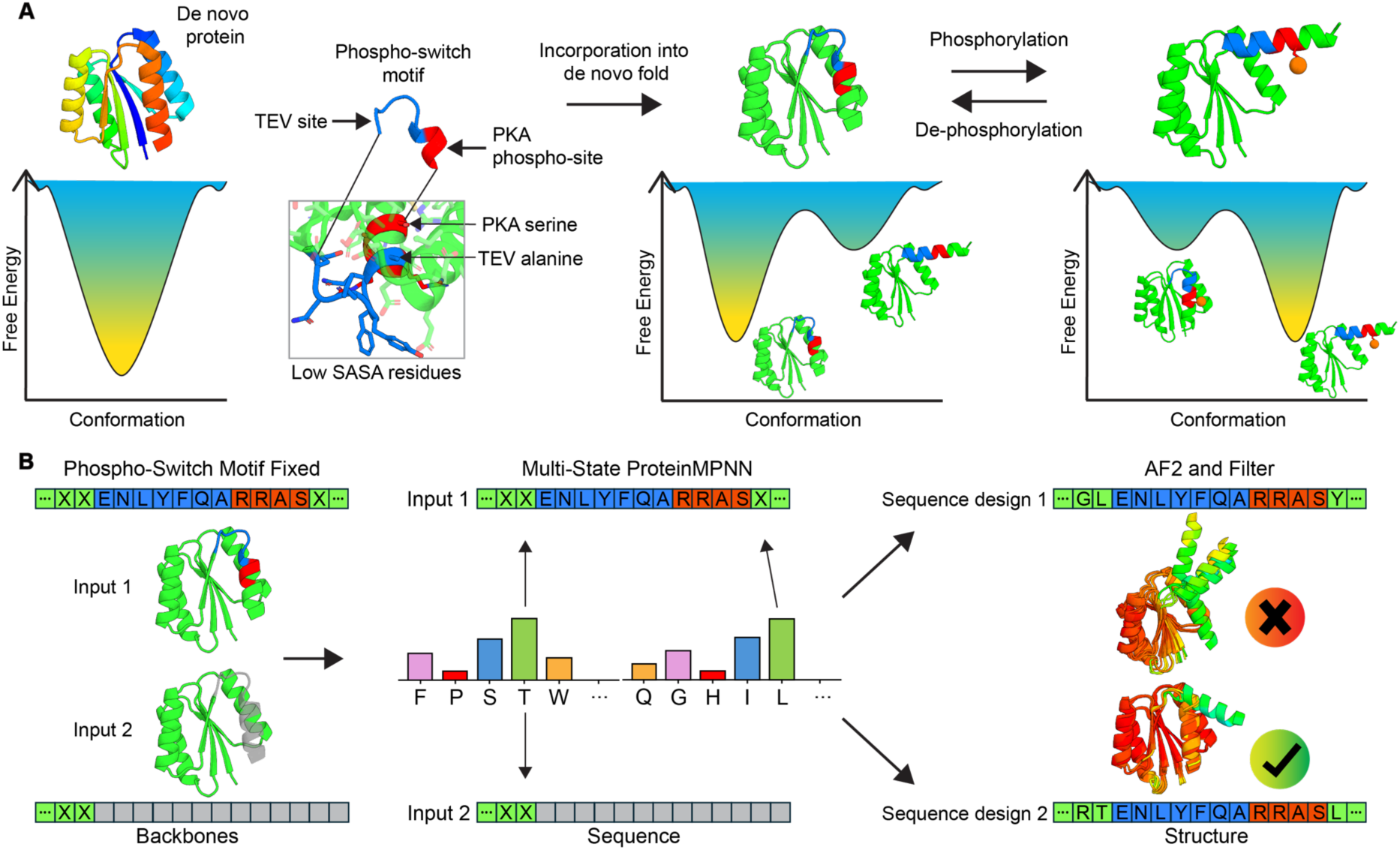
Conceptual overview of the phospho-switch mechanism and computational design approach. **(A)** Schematic representation of the phospho-switch concept. De novo proteins generally adopt a single stable conformation, represented by a single minimum in the energetic landscape. Incorporation of a phospho-switch motif (TEV cleavage site and PKA phosphorylation site) into de novo scaffolds can enable proteins to access two distinct, energetically favorable conformational states. Phosphorylation and de-phosphorylation can regulate equilibrium between the two states. **(B)** Computational design process for state switching proteins. A fixed phospho-switch motif was embedded into de novo scaffolds, such that the phosphorylation site serine and TEV cleavage site were buried in the closed state (input 1). The terminal helix was removed to mimic a phosphorylated/open state (input 2). Multi-state ProteinMPNN used both inputs to generate sequences compatible with both states. Output sequences were evaluated with AF2 and filtered based on confidence and structural metrics.

## Results

### Multi-state design of a phospho-responsive de novo protein

To design state-switching proteins we embedded a phosphorylation site in a buried residue contained within a C-terminal helical segment of several de novo folds. The mechanism we were aiming for within this design was that the phosphorylation would favor the open state (Figure 1A). To monitor the state of the switch we also incorporated a TEV cleavage site that can serve as an orthogonal reporter, meaning that non-phosphorylated proteins are mostly expected to rest in the closed conformation (serine and TEV cleavage site inaccessible) and will be preferentially open in the phosphorylated state, allowing different rates of TEV cleavage for working designs, enabling medium to high-throughput screening workflows.

As a kinase we used PKA, a serine kinase that recognizes the motif RRxS. We used several de novo α/β-sandwich folds with C-terminal helices, as this type of topology enables the shielded placement of the motif’s phosphorylation site serine buried in the core and the recognition arginines exposed to solvent (Figure 1A). We opted to use RRAS as the PKA recognition motif, as alanine facilitates conservation of this region’s helicity in the scaffold proteins. Three known de novo folds with existing structures were utilized: 2LV8^21^, 7BPL^22^, and 5GAJ^23^; referred to as Phosphorylation Induced Protein Switches (PIPS) PIPS1, PIPS2, and PIPS3 respectively. Furthermore, we incorporated a diffusion design pipeline to generate a novel de novo fold, referred to as PIPS4, to interrogate the generalizability of our design and screening approach. For this, random combinations of secondary structural elements (with a fixed C-terminal strand–helix) were fed into Difftopo^24^ to find suitable topologies, which were then refined with RFDiffusion^25^ to generate backbones (Supplementary Figure 1) (see Methods for details).

To design sequences compatible with both states, we employed multi-state ProteinMPNN^26^ to generate single amino acid sequences optimized for the stability of both conformations (Figure 1B). Input 1 was defined as the full-length protein, including the fixed phospho-switch motif, in the closed (non-phosphorylated) conformation. Input 2 had a C-terminal truncation, mimicking the open (phosphorylated) conformation of the protein. The probabilities of residues at each position for each input were combined and sampled to generate sequences capable of occupying both states. C-terminal residues were sampled exclusively from Input 1. Then, AlphaFold2^10^ (AF2) was used to predict the structures of the designed sequences. Designs were filtered based on predicted local distance difference test (pLDDT) and the solvent accessible surface area (SASA) of the phosphorylation site serine and TEV site alanine, to ensure that the closed state shielded the serine and TEV cleavage site to minimize the risk of obtaining a basally open switch and ensuring the functional integrity of the design (see Methods for details).

### Computational phospho-switches are triggered by a kinase

Designs were screened via yeast surface display with a double tag strategy (N-terminal HA and C-terminal V5). The two tags allow us to monitor the display (HA) and the presence of the C-terminal tag (V5). The accessibility of the TEV cleavage site served as a proxy to distinguish closed (inaccessible) from open (accessible) conformations. This approach allowed us to compare the rates of cleavage, measured by loss of V5 signal, between non-phosphorylated and phosphorylated samples to identify designs that cleave faster in a phosphorylation-dependent manner. To validate the functionality of the phospho-switch motif in the context of the screening strategy, 14 PIPS1 designs were tested as single clones, with one design showing phosphorylation-dependent switching. However, after this initial success, the next four scaffolds attempted resulted in no successful switches. The scaffolds 7BQE^22^ and 7BQC^22^ were unable to scaffold the phospho-switch motif while satisfying the SASA filters, and 7BPN^22^ passed the SASA filters, but failed the pLDDT filter. Additionally, the 2LCI^23^ fold passed all in silico filters, but none of the 11 designs tested showed phosphorylation-dependent switching.

Due to these varying results, we sought to assess PIPS2-4 in a higher-throughput manner and screened 253, 762, and 985 designs respectively (Supplementary Figure 2). We devised a screening strategy to identify designs that were sensitive to TEV cleavage upon kinase treatment, while excluding those that were constitutively open. To do so, we employed 3 rounds of screening: first, to remove all the designs that showed TEV cleavage sensitivity without kinase treatment; second, to select designs that were successfully phosphorylated; and third, to select designs that were sensitive to cleavage upon phosphorylation (Supplementary Figure 3). Designs encompassing all the different folds PIPS1-4 (Figure 2A) show switchable characteristics upon phosphorylation, detected by the increase in the rate of TEV cleavage (by decrease in V5 signal) once phosphorylated (Figure 2B,D) (sequences available in Supplementary Table 1). Additionally, phosphorylated designs treated with lambda protein phosphatase (LPP) were able to retain their C-terminal V5 signal, showing the reversibility of the phospho-switch behavior (Figure 2B green histograms). To probe the importance of the phosphorylation site serine designed in the protein core, we tested alanine mutants which lack the capability of being phosphorylated (Figure 2C,E). To confirm that the displayed proteins were successfully phosphorylated, an anti-phosphorylation antibody specific for the phosphorylated PKA motif was used (Figure 2D,E). The validated PIPS1-3 had sequence identities to their initial scaffolds of 35.71%, 37.00%, and 58.59% respectively (Supplementary Figure 4). In summary, we computationally designed using a backbone generation approach coupled to multi-state design and experimentally validated multiple sequences with phosphorylation-dependent switching capabilities at the surface of yeast.

**Figure 2:**
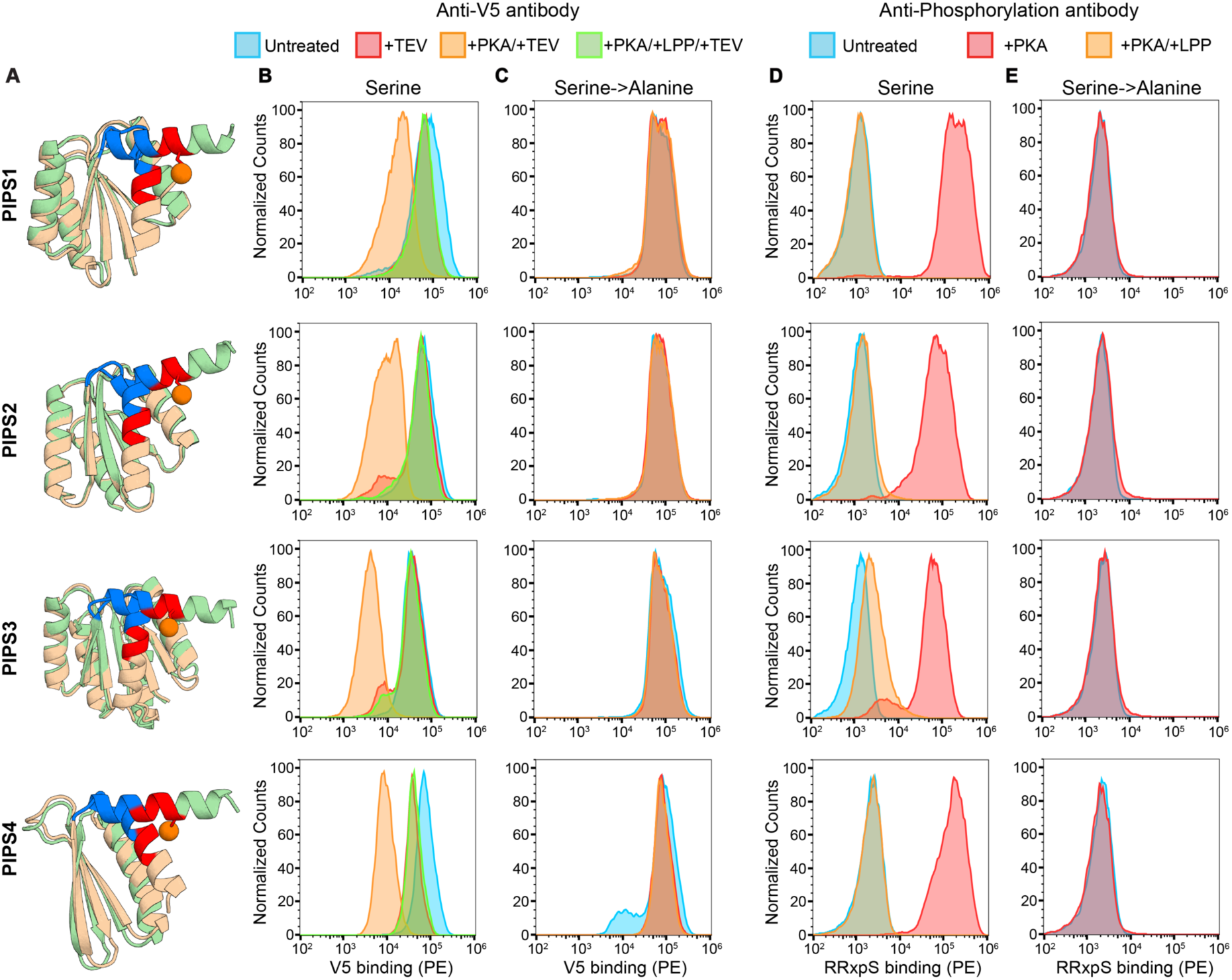
Experimental characterization of phospho-switching de novo proteins. **(A)** Predicted open and closed configurations of PIPS1-4. AF2 generated non-phosphorylated (wheat) and AlphaFold3 (AF3) generated phosphorylated (pale green), with TEV site (blue), PKA motif (red), and phosphate (orange sphere). **(B)** Flow cytometry histograms of PIPS1-4 containing the phosphorylation site serine displayed on yeast, showing binding signal for the C-terminal V5 tag (phycoerythrin, PE) after the indicated treatments. **(C)** Flow cytometry histograms of PIPS1-4 mutants containing a serine to alanine mutation analyzed as in (B). **(D)** Flow cytometry histograms of PIPS1-4 containing the phosphorylation site serine displayed on yeast, showing binding signal for the phosphorylated PKA motif (RRxpS) (phycoerythrin, PE) after the indicated treatments. **(E)** Flow cytometry histograms of PIPS1-4 mutants containing a serine to alanine mutation analyzed as in (D).

### Designed phospho-switches are stable in both open and closed states

PIPS1-4 were expressed, purified, and biochemically characterized (Figure 3). All designs were monomeric in both the non-phosphorylated and phosphorylated states, with molecular weights close to the theoretical values as determined by size-exclusion chromatography coupled to multi-angle light scattering (SEC-MALS) (Figure 3A). All designs were well-folded and thermally stable in both states, as evidenced by circular dichroism (CD) (Figure 3B,C). The phosphorylated state of PIPS4 is the only protein shown to have a T_m_ under 90°C, which is consistent with its smaller size and hydrophobic core. Phosphorylation of the purified PIPS was confirmed by matrix-assisted laser desorption/ionization time of flight (MALDI-TOF) mass spectrometry, where m/z shifts of ∼80 Da (molecular weight of phosphate) were observed for each protein (Figure 3D). The performed biochemical validation showed that the designs are thermally stable and well behaved in both the non-phosphorylated and phosphorylated states, which is a requirement for functional switching proteins.

**Figure 3:**
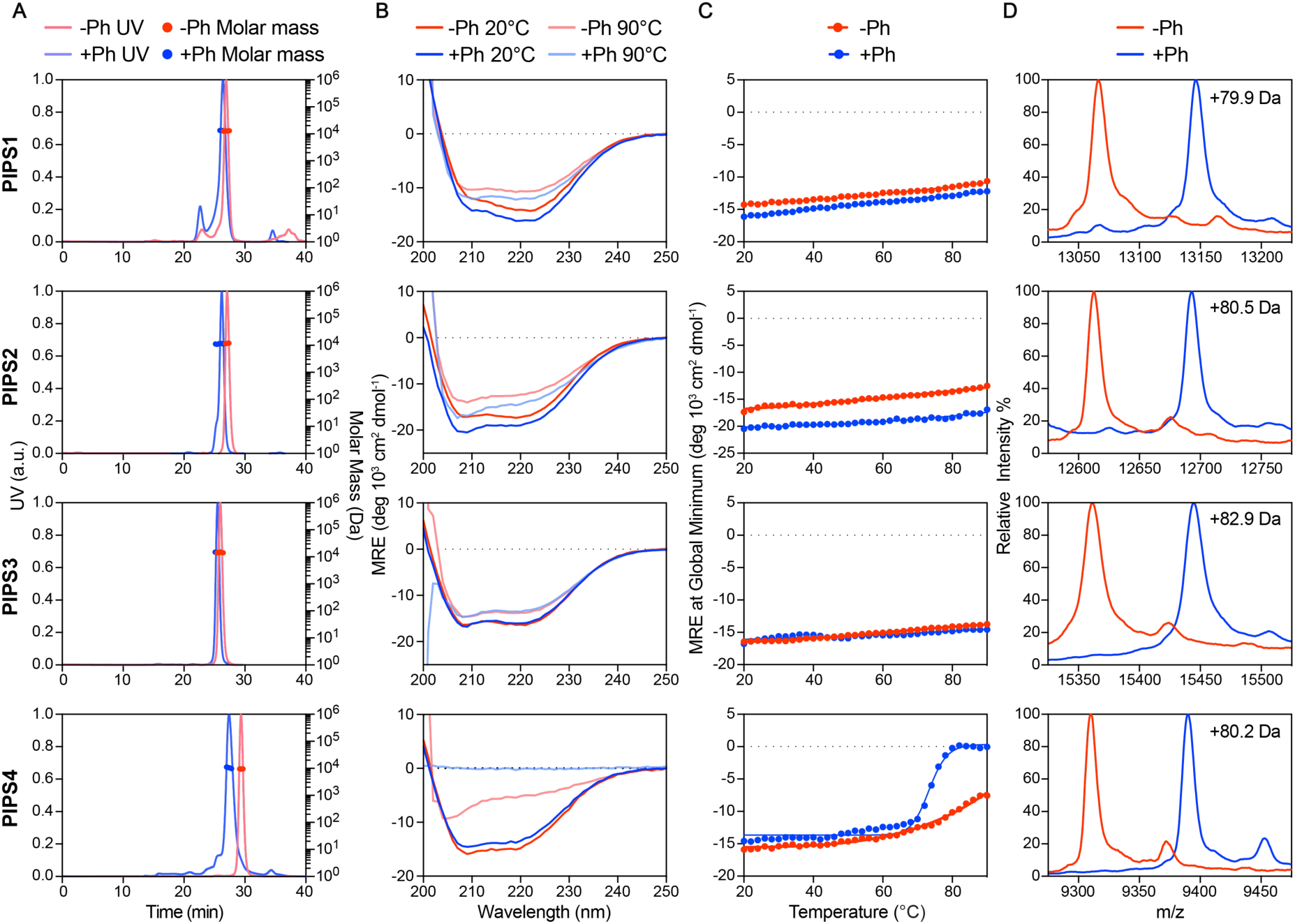
Biophysical characterization of PIPS. **(A)** Size-exclusion chromatography multi-angle light scattering (SEC-MALS) analysis of PIPS in the non-phosphorylated (red) and phosphorylated (blue) states. **(B)** Circular dichroism (CD) spectra of PIPS at 20°C (red, blue) and 90°C (light red, light blue) in the non-phosphorylated and phosphorylated states respectively. **(C)** Thermal denaturation curves derived from CD spectra, plotted by tracking the global minimum of each spectrum between 20°C and 90°C. **(D)** Matrix-assisted laser desorption/ionization time of flight (MALDI-TOF) mass spectra of PIPS1-4 before and after phosphorylation. Increase in m/z between the non-phosphorylated and phosphorylated states of PIPS1-4 are shown in each respective panel.

### PIPS2 structure and dynamics by NMR spectroscopy

To examine the accuracy of the designs and explore how structure and dynamics allow them to control the switching while limiting the population of the open conformation in the non-phosphorylated state, we studied the PIPS2 design by solution-state NMR spectroscopy. Structure determination of the non-phosphorylated state revealed a very accurate design, with a Cα root mean square deviation (RMSD) of 1.70Å between the computational model and the determined structure (model 1 of the NMR ensemble) with both serine-96 (S96) (the designed phosphorylation site serine) and the TEV cleavage site in a shielded configuration (Figure 4A) (structure statistics in Supplementary Table 2). 15% of the residues lack backbone ^1^H,^15^N assignments, mapping to Thr85-Arg94 (just before S96) in addition to Lys41, His42, Tyr43 and Lys60 which are all packed against the Thr85-Arg94 segment in the 3D structure (Supplementary Figure 5). This whole region displays a high Cα root mean square fluctuation (CA RMSF) in the NMR ensemble (Figure 4B, Supplementary Figure 6), which together with the missing ^1^H,^15^N resonances hints at a whole region experiencing extensive dynamics around the micro-to millisecond timescale^27,28,29^. We then studied PIPS2’s structural dynamics in more detail through ^15^N relaxation and solvent exchange experiments. A set of residues (Val20, Glu38, Val47, Leu54, Ile55, Arg62, Leu84) display ^15^N R_2_/R_1_ values above those expected for the central value that corresponds to a global correlation time of 7.2 ns consistent with monomeric protein. Another set of residues (Ala4, Asp5, Ile7, Asp15, Ile36, Val39, Met44-Ser48, Gly52, Leu56, Ala57, Glu66, Ala68, Glu70-Glu75, Asp78-Glu82, Ala95) display shifts for excited states in ^15^N CEST experiments (Figure 4C, Supplementary Figure 7).

**Figure 4:**
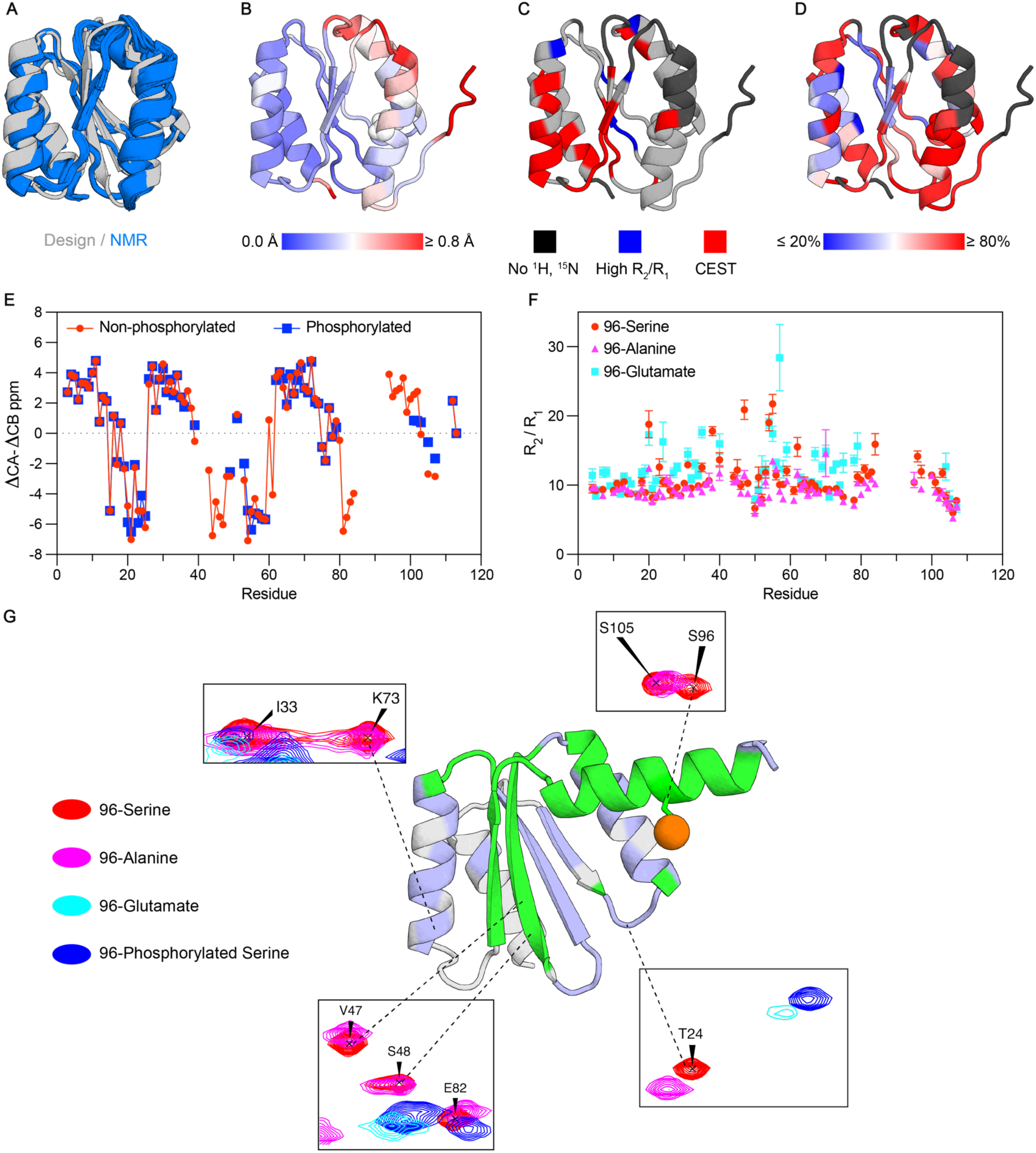
Structural analysis of PIPS2. **(A)** Superimposition of original AF2 model for PIPS2 (gray) and the NMR ensemble for the determined NMR structure (blue). **(B)** Model 1 from the deposited NMR structure shown as cartoons with CA RMSF mapped in a scale from blue = 0 Å to red ≥ 0.8 Å, with S96’s sidechain shown as spheres. **(C)** Model 1 from the deposited NMR structure shown as cartoons colored by the different cues for dynamics (black = missing data, blue = high R_2_/R_1_ ratio, red = excited state peak in ^15^N CEST). **(D)** Model 1 from the deposited NMR structure shown as cartoons colored by the fraction of intensity lost in the crosspeaks of a ^1^H,^15^N HSQC spectrum after 25 minutes of exchange with buffer prepared in ^2^H_2_O. Blue: ≤ 20% of signal lost, red: ≥ 80% of signal lost. Higher fraction of signal lost is indicative of higher solvent exposure. **(E)** Differences in ^13^C CA and CB chemical shifts relative to the values expected for the corresponding residues in fully disordered states (ΔCA-ΔCB) for the non-phosphorylated (red) and phosphorylated (blue) states. Positive ΔCA-ΔCB values indicate helical secondary structures and negative values indicate sheet conformations. **(F)** R_2_/R_1_ ratios for PIPS2 with a serine, alanine, or glutamate at position 96. **(G)** AF3 model of phosphoPIPS2 with phosphate on serine-96 shown as an orange sphere and NMR data for the phosphorylated form mapped as follows: green = missing crosspeaks, light blue = large ^1^H,^15^N chemical shifts, gray = similar to non-phosphorylated PIPS2. Insets: portions of ^1^H-^15^N HSQC spectra of non-phosphorylated PIPS2 (red), S96A mutant (pink), S96E mutant (cyan), and phosphoPIPS2 (blue).

Globally, these observations demonstrate a hierarchy of timescales in dynamics around the designed switch (Figure 4C), spanning from around 10-1,000 1/s (crosspeaks giving a CEST response) to 100-5,000 1/s (crosspeaks broadened beyond detection) and 1000-10,000 1/s (crosspeaks visible and with high R_2_/R_1_). Fits of the ^15^N CEST data to a two-state Bloch-McConnell model for chemical exchange show excited states populated by around 2% and exchanging with the ground state at around 50-100 1/s (Supplementary Figure 7). Deuterium exchange experiments show that PIPS2’s interior is quite accessible to the solvent despite its overall folded nature (Figure 4D). In particular, the backbone HN protons in and around the region that contains the designed switch exchange faster than those in the other helices, even though the region is structured overall and the ^13^C assignments show a well-defined helix (Figure 4E, red trace). These results suggest that the PIPS2 design samples breathing motions that allow for solvent exchange at its interior during incursions to excited states with a substantial degree of exposure as required to allow for phosphorylation.

On analysing phosphorylated PIPS2 (referred to as phosphoPIPS2) by NMR, we observed that crosspeaks in the ^1^H,^15^N HSQC were more heterogeneous in intensity than in the non-phosphorylated species, some even too broad to be assigned (Supplementary Figure 8). We found large ^1^H,^15^N chemical shift perturbations relative to non-phosphorylated PIPS2 (Supplementary Figure 9) and 14 additional residues lacking assignments (same as for non-phosphorylated PIPS2 plus Asp61, Arg62, Gln80-Leu84 and Ala95-Glu101). However, the ^13^C chemical shifts for the residues assigned in both species were very similar, producing almost identical ΔCA-ΔCB patterns (Figure 4E). These observations suggest that the global fold in phosphoPIPS2 is likely similar to that of the non-phosphorylated form, but the dynamics already present in the non-phosphorylated form are more pronounced, affecting the protein to a larger extent in terms of number of residues involved and/or the extent of dynamic open states.

To further probe the PIPS2 design, we characterized and studied mutants of S96 to alanine (S96A) and glutamate (S96E), corresponding to constitutively closed and open states, respectively (Supplementary Figure 10). The S96A mutant displayed an HSQC spectrum very similar to that of S96-PIPS2 (Supplementary Figure 8), and while it did not recover crosspeaks that could be assigned to those missing in S96-PIPS2 and CEST experiments still revealed 2% population of an excited state, this mutant did not present any residues with ^15^N R_2_/R_1_ above the value expected for the global correlation time of the monomer, meaning that only dynamics in the faster end of the slow timescale are quenched in this mutant (Figure 4F). On the contrary, the S96E mutant displayed an HSQC spectrum closer to that of phosphoPIPS2, also presenting heterogeneous intensities; in this case, ^15^N CEST detects no residues undergoing slow timescale dynamics, while ^15^N R_2_/R_1_ profiles show a large number of residues with values that indicate dynamics in the faster end of the slow-timescale regime (Figure 4F). We interpret this as the S96E mutant, and by analogy the phosphoPIPS2, being less structured around the engineered positions, with the C-terminal helix possibly exchanging among multiple open conformations on the faster end of the slow timescale while the core of the fold remains overall intact (Figure 4G).

### Design of conformation-specific binders

Conformation-specific PPIs enable conditional interactions that can control the responsiveness of signaling processes. To generate a second effector that could read the state of a designed phospho-switch and enable a phospho-dependent protein interaction, we sought to design binders specific to the phosphorylated state of PIPS2. We designed binders targeting phosphoPIPS2 using two different approaches, BindCraft^30^ and a Rosetta MotifGrafting approach^31^ (Supplementary Figure 11). In both approaches we targeted the site unveiled by the open conformation of PIPS2. The MotifGrafting approach used the terminal helix motif in the PIPS2 design as the interaction mediator, and this motif was grafted onto scaffold proteins (see Methods for more details).

The designed binders were screened by yeast display (Supplementary Figure 12) using both PIPS2 and phosphoPIPS2, the best designs were identified using a deep sequencing read out. Selected designs were screened as single clones and two binders of interest, referred to as PIPS2-B1 and PIPS2-B2, were confirmed to be highly specific towards phosphoPIPS2 (Figure 5B,C). Additionally, interface point mutants of these binders were tested and shown to completely abrogate binding, suggesting the designed interfaces are accurate.

**Figure 5:**
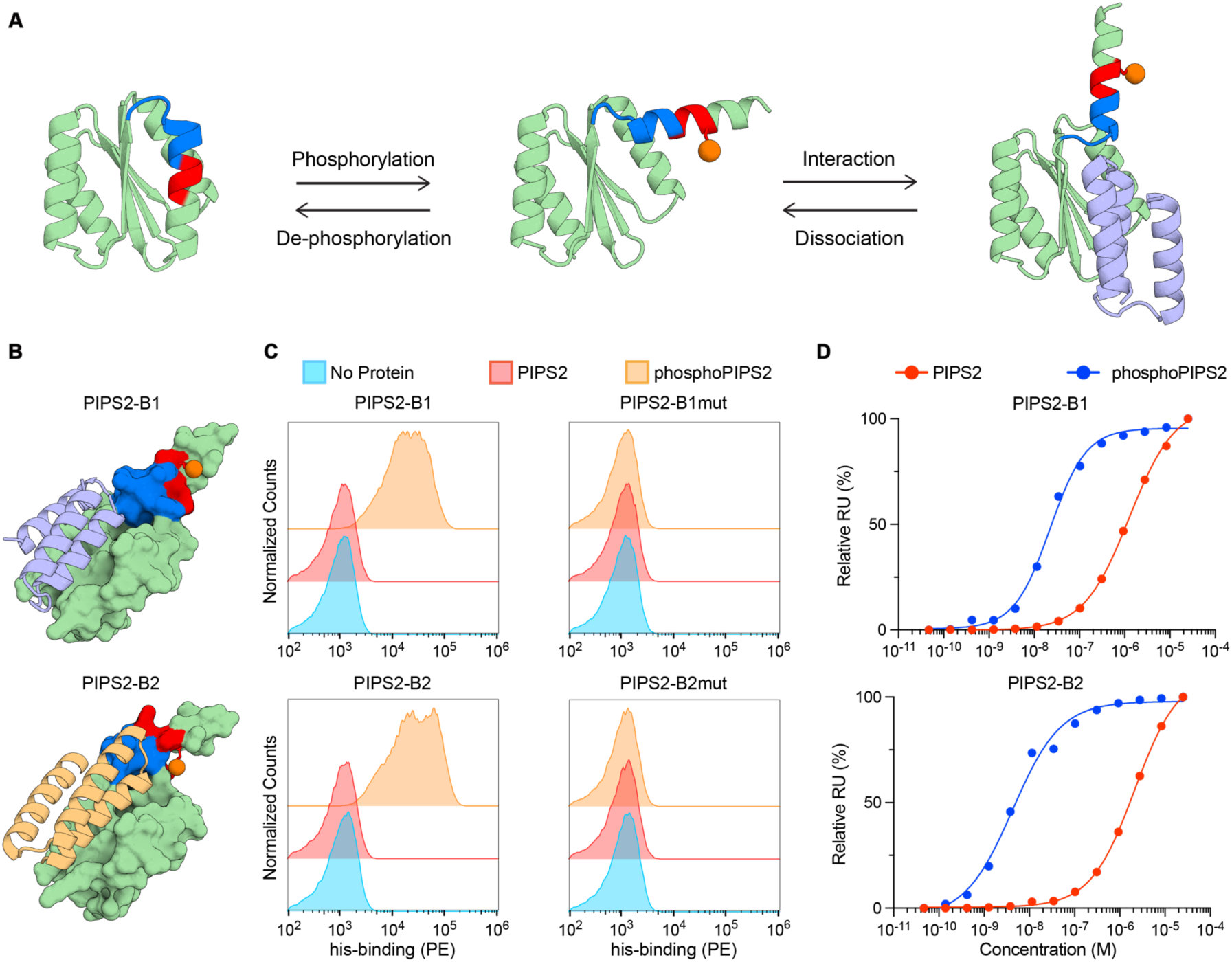
Biochemical characterization of conformation-specific binders to phosphoPIPS2. **(A)** Schematic representation of the phosphorylation-dependent binding mechanism of PIPS2. Left, PIPS2 predominantly occupies a closed conformation. Middle, upon phosphorylation phosphoPIPS2 is driven towards an open conformation. Right, the open conformation exposes an interface to facilitate a PPI. **(B)** AF3 multimer models of designed binders PIPS2-B1 (light blue) and B2 (light orange) bound to phosphoPIPS2 (pale green) shown as a surface representation. For PIPS2, blue represents the TEV site, red represents the PKA site, and the orange sphere represents the phosphate. **(C)** Flow cytometry histograms showing binding signal (phycoerythrin; PE) of single clone yeast displaying designed binders PIPS2-B1 and PIPS2-B2. Yeast were either incubated without or with 100nM of the respective target protein, PIPS2 or phosphoPIPS2. Binder interface mutations of PIPS2-B1mut (A23R) and PIPS2-B2mut (A53R) were used as specificity controls. **(D)** Normalized steady-state SPR data of PIPS2-B1 and B2 against PIPS2 and phosphoPIPS2. Kds were calculated by fitting a nonlinear four-parameter curve to the data.

Next, the binders were purified and characterized (Supplementary Figure 13) and surface plasmon resonance (SPR) was conducted to measure their affinity to both PIPS2 and phosphoPIPS2 (Figure 5D). PIPS2-B1 and PIPS2-B2 have dissociation constants (Kds) of 22.39 nM and 4.28 nM to the phosphoPIPS2, respectively. A Kd could not be accurately determined for the non-phosphorylated state of PIPS2 for either binder, however, the values exceed 1 µM, showing that the binders have a preference for the phosphorylated state by multiple orders of magnitude.

### Assembling minimal signaling networks in cell-based systems

The relay of biochemical signals is at the heart of virtually all biological processes. The ability to rationally engineer protein modules to build such signaling systems from scratch could bring insight to our understanding of native pathways as well as for the creation of novel cellular pathways.

To test PIPS2’s ability to act as a phosphorylation-based switch in cells, a split transcription factor system was utilized, where PIPS2 was fused to the DNA binding domain Gal4 and the validated binders were fused to the transcription activation domain RelA (Figure 6A). Here, we hypothesized that the phosphorylation-dependent interaction could recruit RelA to the Gal4, reconstituting a functional transcription factor that initiates transcription of a gene for the reporter protein secreted NanoLuciferase^32^. This simplistic signaling network approach mimics natural signaling cascades, where the proximity induced activation of transcription is regulated by the phosphorylation of PIPS2. Endogenous PKA was activated by addition of the small molecule forskolin, which increases intracellular cAMP, thereby releasing the regulatory inhibiting domains of PKA and freeing the catalytic domain^33^. We show that 25 µM of forskolin induces approximately a 4-fold increase for both PIPS2-B1 and B2 (Figure 6B).

**Figure 6:**
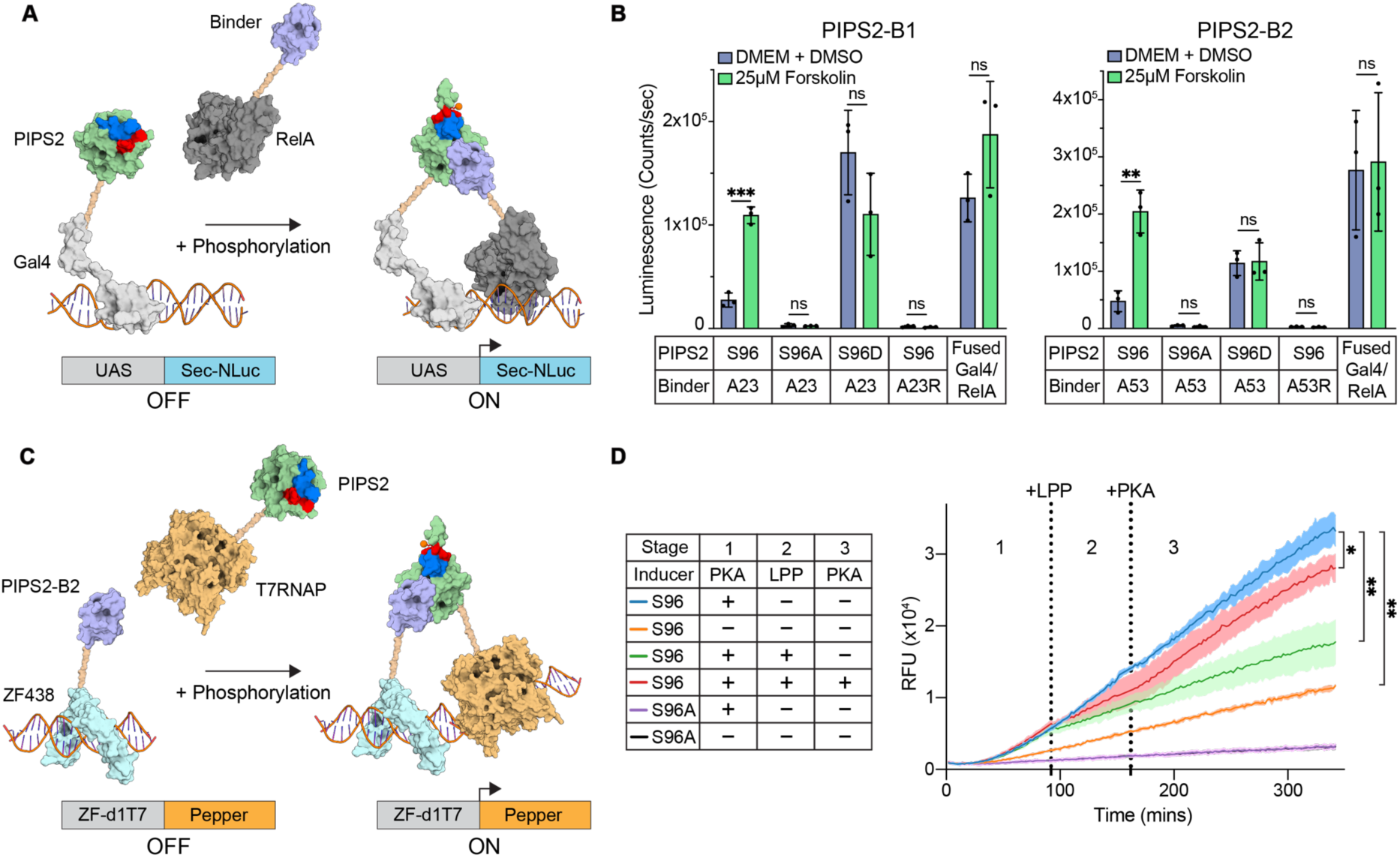
PIPS2 is functional in cell-based systems. **(A)** Schematic of the split transcription factor assay where PIPS2 is fused to a Gal4 DNA binding domain and binders are fused to a RelA transcription activation domain. Phosphorylation-induced interactions between PIPS2 and the binder drives transcription of a UAS (upstream activating sequence)-driven secreted NanoLuciferase reporter (Sec-NLuc). **(B)** Luminescence measurements for HEK293T cells transfected with PIPS2-Gal4 and either (left) PIPS2-B1-RelA or (right) PIPS2-B2-RelA. Point mutants of both PIPS2 (S96A, S96D) and binders PIPS2-B1 (A23R) and PIPS2-B2 (A53R) were used as controls. Cells were either untreated or induced with 25µM forskolin. A fused Gal4-RelA construct served as a positive control. **(C)** Schematic of a cell-free expression system where PIPS2 is fused to T7 RNA polymerase and PIPS2-B2 fused to a DNA binding zinc finger (ZF438). Phosphorylation-induced interactions between PIPS2 and binder recruits T7 RNA polymerase to activate transcription of the fluorescent Pepper aptamer reporter gene. **(D)** Fluorescence measurements from a three-stage cell-free transcription assay. Left, table indicating PIPS2 variant used in each group and their respective treatments with (+) or without (-) the indicated inducer at each stage. Right, line plot of measured fluorescence of Pepper aptamer over time. Time points for addition of LPP (Stage 2) and PKA (Stage 3) shown as a vertical dotted lines. Data in graphs **(B)** and **(D)** represent mean +/-s.d. Each assay was performed using n = 3 biological replicates. Statistical significance was determined using unpaired two-tailed Welch’s t-tests (ns = not significant, * = p < 0.05, ** = p < 0.01, *** = p < 0.001).

Additionally, we used mutants of both PIPS2 and the binders to test the role the phospho-switch plays in signal induction. Two mutants were utilized for PIPS2, S96 was mutated to alanine (S96A) and aspartate (S96D). The alanine mutant mimics a constitutively closed form of PIPS2, while the aspartate acts as a phospho-mimetic to mimic a constitutively open state. As predicted, the alanine mutant showed low response, while the aspartate mutant enhanced reporter expression, evidencing the role of the phospho-controlled conformational change in regulating the binder’s access to the interface. The split transcription factor components were fused into a single chain (fused Gal4/RelA) to serve as a control to monitor general protein expression of cells treated with forskolin. The only samples that show a significant increase in signal upon forskolin induction were the non-mutated PIPS2 in combination with the native binders. The fused Gal4-RelA construct did not significantly increase signal, suggesting the increase in signal from the PIPS2/binder pairs is not due to a general increase in protein expression. Additional controls were included to validate assay conditions. The CREB transcription factor, which is directly activated by PKA, was used to monitor endogenous PKA activity via a CREB-activated reporter plasmid expressing secreted NanoLuciferase. As a negative control, H89, a competitive inhibitor of ATP, was used for its well characterized inhibitory effects on PKA^34^ (Supplementary Figure 14).

To further investigate whether the increase in signal is due to the phosphorylation-dependent PPI, a cell-free assay was conducted in a more controlled environment with fixed amounts of protein based on a reported platform^35^ (Figure 6C). When PKA was added to the reaction, the fluorescence signal from the Pepper aptamer reporter increased, showing that PIPS2 can regulate transcription in a phosphorylation-dependent manner (Figure 6D). At the timepoint of 82 minutes, the indicated samples were spiked with LPP (green and red), while the other samples received only LPP buffer. Then, at the timepoint of 152 minutes, the indicated sample (red) was spiked with additional PKA, while the other samples received PKA buffer only. This data shows that when LPP is added to the ongoing reaction, the transcription rate decreases to baseline. Furthermore, when additional PKA is added to the reaction, we again see an increase in transcription by PKA phosphorylation outcompeting the dephosphorylation rate of LPP. Overall, this data suggests that the designed proteins can be assembled into a minimal, reversible signaling network, which we employed to regulate transcription in a phosphorylation-dependent manner.

## Discussion

While many natural proteins utilize PTMs to transition between non-functional and functional states, de novo protein design has so far mostly focused on stable single state proteins. Designer proteins that switch between two states generally need a stimulus that will shift their energetic landscape towards a different energetically favorable state. Here, we aimed to extend de novo protein design to proteins capable of occupying two distinct states regulated by phosphorylation. By shielding the serine of the PKA recognition motif and a TEV cleavage site in the closed state, we were able to validate the phosphorylation-induced switchability of four folds (PIPS1-4).

Similar to natural signaling proteins, these PIPS sample different conformations, with the population of the dominant state regulated by phosphorylation in a reversible manner. In the phosphorylated state, the open conformation becomes predominant, and NMR data for phosphoPIPS2 indicates the global fold is maintained in this state (Figure 4E). This structural conservation enabled the design of binders that target an interface that is only accessible in the open conformation, thereby regulating the interaction in a phosphorylation-dependent manner. These binders allowed us to create a simple synthetic signaling cascade capable of regulating transcription in response to PKA activity. While this study focused primarily on Rossman-like folds, we speculate that this approach could be expanded to more diverse protein topologies by incorporating other protease cleavage sites and/or kinase recognition motifs.

The combination of the multi-state design approach coupled with the AF2-based filtering allowed for the selection of designs capable of phosphorylation-induced switching while retaining their ability to remain well-folded and monomeric in both non-phosphorylated and phosphorylated states. This approach’s ability to generate and select sequences that are compatible with both states is most evident in PIPS3, which has a glutamate in its core in the non-phosphorylated state (Supplementary Figure 15). This glutamate is predicted to form a stabilizing h-bond network with the phosphorylation site serine and a nearby threonine in the closed conformation, while acting as a polar surface residue in the open conformation. Additionally, the fact that the S96A PIPS2 mutant retains dynamics on the slower range of the monitored timescales shows that the design and selection process requires more than simply placing a shielded serine in the core; it must also tune the dynamics required to eventually allow transient exposure of the serine. Experiments, modeling, and molecular dynamics simulations have shown that natural proteins often expose hidden phosphorylation sites in their non-phosphorylated states as they undergo local and global dynamics, covering a wide range of timescales from picoseconds to seconds^17,36,37^. This balance between a predominantly closed conformation in the non-phosphorylated state while retaining a minor population that remains accessible to the kinase at equilibrium is paramount to the design of successful phospho-switches.

The ability of AF3^11^ to model proteins with phosphorylated residues offers promise for such design tasks. However, it should be noted that differences between AF2 and AF3 predictions exist to varying extents. For example, AF3 models of PIPS1 and PIPS3 would have failed the in silico filters used (Supplementary Figure 16). Additionally, AF3-multimer could not effectively differentiate the binding propensity of PIPS2-B1 and B2 between the non-phosphorylated and phosphorylated states of PIPS2, as all 5 output models were shown to be bound even for the non-phosphorylated PIPS2 target (Supplementary Figure 17).

Developing PIPS that respond to different kinases could lay the foundation for the development of cellular logic gates, incorporating multiple PIPS which respond to different kinases. The design of de novo signaling proteins could be extended to tyrosine kinases, to transduce signals in T cells. Incorporation of such switches into CAR-T cell receptors would allow for the design of AND, OR, and NOT cellular logic gates, regulating CAR-T cell’s activity in an autonomous manner. Additionally, the design of novel kinase/substrate pairs would be advantageous, as it would obviate the reliance on endogenous signaling pathways, mitigating potential activation of off-target pathways. In combination with modular synthetic receptor platforms^38^, we envisage that a synthetic signaling pathway that operates orthogonally to endogenous pathways could be achieved.

Altogether, we show a novel approach for designing and screening de novo proteins that change conformation in a phosphorylation-dependent manner. Together with the phosphorylation-dependent binders, we show how de novo proteins can be used to control kinase dependent signaling cascades to express a target gene. This concept could be extended to orthogonal kinases, providing a blueprint for designing and screening proteins capable of functioning as components in cellular logic gates for cell-based therapeutics or synthetic biology tools.

## Methods

### Selection of scaffolds for hosting phosphorylation sites

To design a protein conformational switch without disrupting the overall stability of the protein, the conformational change should be localized to a terminal region, thereby minimizing the potential for global destabilization. To identify scaffolds that can host a phosphorylation site while maintaining a stable “closed” conformation, we selected protein folds featuring a C-terminal helix. For these folds, we identified amino acid positions at N and N+4 with a combined solvent-accessible surface area (SASA) of less than 3 Å², enabling the transplantation of both the TEV protease cleavage site (ENLYFQ/A) and the PKA phosphorylation site (RRAS). In contrast to the canonical TEV cleavage site of Q/S, we used Q/A as it remains efficiently cleaved by TEV and preserves helicity permitting the TEV cleavage site to be shielded. We explored a diverse set of scaffolds, including pre-existing de novo α/β-protein folds, as well as novel de novo folds generated using the DiffTopo^24^ and RFdiffusion^39^ pipelines. These novel folds were developed to test the generalizability of our approach, where we generated de novo scaffolds with C-terminal helices. As Supplementary Figure 1 shows, we first randomly sampled secondary structure compositions of length 5–7, enforcing a helix at the C-terminus, as input for DiffTopo to produce initial blueprints. These blueprints were then refined via RFdiffusion’s motif scaffolding and partial diffusion pipeline to improve designability, while preserving the global secondary structure stacking. This process yielded hundreds of backbones with high design confidence (ProteinMPNN designs scRMSD < 3, AlphaFold2 pLDDT > 90). After applying the same SASA-based filtering, to ensure compatibility with the target phosphorylation site, we selected 9 final scaffolds for experimental validation in the end. The pre-existing de novo folds included PDB structures 2LV8, 7BPL, and 5GAJ (PIPS1-3 respectively). To ensure PIPS2 and PIPS3 designs remained within the oligo pool gene synthesis size limit (300bp), their N-terminal regions were fixed and built into their respective insertion plasmid, while the rest of the protein was redesigned.

### Customized multi-state ProteinMPNN design for dual-state compatibility

To design a sequence compatible with both the closed and open states, we developed a customized multi-state ProteinMPNN algorithm. Unlike the original multi-state ProteinMPNN, our approach accommodates input backbones of differing lengths. Specifically, we provided two chains as input: the full-length protein, including the C-terminal helix (representing the “closed” state) and a truncated protein, lacking the C-terminal helix (representing the “open” state). We combined the residue probabilities at corresponding positions across the two states, however for residues in the C-terminal helix, only the “closed” state probabilities were considered. Amino acid sequences were then sampled based on the averaged probabilities while keeping the TEV and phosphorylation sites fixed.

### Computational filtering

After generating sequences using ProteinMPNN, we implemented a multi-step computational filtering process. First, to mitigate off-target signals during screening, we ensured that the designed sequences contained no additional phosphorylation motifs beyond the target site, excluding sequences with RRxS, RRxT, KKxS, KKxT, KRxS, KRxT, RKxS, or RKxT motifs. Next, we predicted the structures of the designed sequences using ColabFold^40^ in single sequence mode and applied the following filtering criteria: overall predicted local distance difference test (pLDDT) scores >75 for >3 models, and RMSDs between “closed” states <3.0 Å. Additionally, models for designs were subjected to filtering based on their predicted conformational propensity to be closed or open. The closed/open state was determined using helix alignment gaps and a SASA threshold. For the helix alignment gaps, the designed backbone’s C-terminal helix was aligned to the predicted structure using TM-align^41^. If there were fewer than 5 gaps in this helix-to-helix alignment, the model was considered to be closed, while 5 or more gaps was considered open. For the SASA threshold, a closed conformation must have a solvent-accessible surface area (SASA) <5 Å² for both the phosphorylation site serine and the TEV site’s alanine, a more lenient SASA filter of <10 Å² was originally used for PIPS1 designs. If the helix alignment gaps were 5 or higher, the conformation was considered open and no further SASA filter was required. Designs with ≤ 3 open states passed the filter. This filtering ensured a range of closed and open state predictions to facilitate the screening of a variety of proteins.

### Yeast display library screening

Each library consisted of 2000 designs that were sequence optimized for *Saccharomyces cerevisiae* and ordered as oligo pools from Twist Biosciences. Each gene contained identical 5’ and 3’ sequences to allow for amplification via PCR. A second PCR was performed to add on homology regions to their respective insert vector. Amplified oligo pools and their respective linearized pCTcon2 vectors were transformed at a 5:1 ratio into EBY-100 yeast via electroporation as previously described^42^. Transformation efficiencies were at least 10-fold the size of their respective library. Transformed yeast were grown at 30°C and passaged at least once with minimal glucose medium (SDCAA) before being induced with minimal galactose medium (SGCAA) overnight. The next morning, yeast were centrifuged for 3 minutes at 2500xg and washed with 0.1% BSA/PBS before beginning experiments. Samples were aliquoted into their respectively labelled tubes, centrifuged again, and supernatant discarded, allowing them to be resuspended in the appropriate buffer conditions for the respective experiment. All fully processed samples were washed and resuspended in appropriate volumes of 0.1% BSA/PBS before being analyzed. All samples were processed and sorted using the Sony SH800 cell sorter (LE-SH800SZFCPL Cell Sorter, v.2.1.5) and data was analyzed using FlowJo (BD Biosciences, v.10.8.1).

### Fluorescence activated cell sorting of PIPS

The amplified genes for the PIPS2-4 were transformed together with their respective pCTcon2 digested vectors containing their appropriate homology regions. All vectors contained a N-terminal HA-tag and a C-terminal V5-tag. Three different sorts were performed to filter designs that contain phosphorylation-induced switching behavior.

#### Sort 1: Aim – To get rid of constitutively open-state proteins

The pre-washed PIPS2 and PIPS4 samples were resuspended with 2µM SuperTEV^43^ in TEV buffer (20mM Tris, 150mM NaCl, pH = 7.5), and the pre-washed PIPS3 sample was resuspended with 15µM SuperTEV. After a 1 hour incubation for PIPS2 and PIPS4 and 30 minute incubation for PIPS3 at 30°C, cells were washed and labelled with the primary antibody mouse anti-V5 (MA515253, Invitrogen;1:300 dilution) for 30 min at 4°C. Cells were washed and labelled with FITC-conjugated goat anti-HA tag antibody (Bethyl; ref: A190-138F; display tag; 1:100 dilution) and PE-conjugated goat anti-mouse antibody (Invitrogen; 12-4010-82; 1:200 dilution). Yeast that retained a PE signal were sorted.

#### Sort 2: Aim – To select proteins that get phosphorylated

Yeast were resuspended either with or without 600nM PKA in PKA buffer (50mM Tris-HCL, 10mM MgCl_2_, 0.1mM EDTA, at pH 7.5) with 5mM ATP and incubated shaking at 30°C for 1 hour. Samples were washed and labelled with the primary antibody rabbit anti-Phospho-PKA Substrate (RRXS*/T*) (9624S, Cell Signaling; 1:200 dilution) for 30 min at 4°C. Cells were washed and labelled with FITC-conjugated goat anti-HA tag antibody (Bethyl; ref: A190-138F; display tag; 1:100 dilution) and PE-conjugated donkey anti-rabbit antibody (406421, BioLegend; 1:160 dilution) for 30 min at 4°C. Yeast that showed a PE signal were sorted.

#### Sort 3: Aim – To select proteins that exhibit phosphorylation induced switching behavior

All samples were resuspended with +Phosphorylation (600nM PKA) buffer and incubated shaking at 30°C for 1 hour. Cells were washed and the PIPS2 and PIPS4 samples were resuspended in a buffer either with or without 2µM SuperTEV in TEV buffer, the PIPS3 samples were resuspended either with or without 15µM SuperTEV. After a 1 hour incubation for the PIPS2 and PIPS4 samples and 30 minute incubation for the PIPS3 samples at 30°C, cells were washed and labelled with the primary antibody rabbit anti-Phospho-PKA Substrate (RRXS*/T*) (as described above) for 30 min at 4°C. Cells were washed and labelled with FITC-conjugated goat anti-HA tag antibody (as described above) and PE-conjugated donkey anti-mouse PE (as described above) for 30 min at 4°C. Phosphorylated yeast that showed a decrease in PE signal upon TEV treatment were sorted. Sorted yeast were plated on SDCAA agar plates. Single colonies were picked, sequenced, and tested for switchable behavior as indicated below in the ‘Single clone PIPS validation via yeast surface display’ section.

### Single clone analysis via yeast surface display

All single clone yeast samples were prepped as follows. Genes encoding single designs were obtained from Twist Bioscience, containing homology overhangs for cloning. Purified and sequence-verified plasmids were transformed into competent EBY-100 yeast using the Frozen-EZ Yeast Transformation II Kit (Zymo Research). Transformed yeast were grown in minimal glucose medium (SDCAA) +P/S medium at 30°C and passaged once before being induced with minimal galactose medium (SGCAA) medium overnight. The next morning, yeast were centrifuged for 3 minutes at 2500xg and washed with 0.1%BSA/PBS before beginning experiments. All single clone analysis was performed on a Gallios flow cytometer (Beckman Coulter) and data was analyzed using FlowJo (BD Biosciences, v.10.8.1).

### Single clone PIPS validation via yeast surface display

Each design was cloned into a pCTcon2 plasmid (with a N-terminal HA-tag and a C-terminal V5-tag). PIPS1 was exclusively tested as single clones and thus not involved in the oligopool screened. Induced yeast were washed, aliquoted to their respective tubes, centrifuged again, and supernatant was discarded. Yeast were then resuspended either with or without 600nM PKA in PKA buffer (as described above) with 5mM ATP incubated shaking at 30°C for 1 hour. After washing the cells, samples were either resuspended in buffer with or without SuperTEV (PIPS1: 2µM, PIPS2: 2µM, PIPS3: 10µM, PIPS4: 15µM) in TEV buffer and incubated shaking at 30°C. After one hour, cells were washed and resuspended either with or without 400 units of lambda protein phosphatase (P0753S, NEB) in 50mM Hepes, 100mM NaCl at pH 7.5 and incubated shaking at 30°C. After one hour, cells were washed and labelled with the primary antibody mouse anti-V5 (as described above) for 30 min at 4°C. Cells were washed and labelled with FITC-conjugated goat anti-HA tag antibody (as described above) and PE-conjugated goat anti-mouse antibody (as described above). For the phosphorylation controls, samples were labelled with the primary antibody rabbit anti-Phospho-PKA Substrate (RRXS*/T*) (as described above) for 30 min at 4°C. Cells were washed and labelled with FITC-conjugated goat anti-HA tag antibody (as described above) and PE-conjugated donkey anti-mouse PE (as described above) for 30 min at 4°C. Cells were washed, and resuspended in appropriate volumes of 0.1% BSA/PBS before being analyzed.

### Structure predictions using AlphaFold3

The same sequences from the original PIPS1-4 designs that underwent AF2 filtering were predicted with AF3^11^ both without and with the phosphorylation site serine phosphorylated. The predictions were run with the default settings and without multiple sequence alignments (MSAs) to compare to the single sequence mode used in AF2. AF3-multimer was used to predict the binding of the PIPS2 binders to PIPS2 in both the non-phosphorylated and phosphorylated states, using the same settings as above.

### Designing binders to phosphoPIPS2

Binders to phosphoPIPS2 were designed using two binder design pipelines, BindCraft and a MotifGraft approach followed by ProteinMPNN and AF2. As a starting point, we decided to use an AF2 PIPS2 model with S96 to alanine mutation to mimic a ‘bound’ conformation. Then the terminal loop and helix was removed, starting from residue E86, allowing us to use the remaining protein segment as the context (target) representing phosphoPIPS2 where the terminal helix is displaced. For BindCraft, this truncated model was used as the target, with residues leucine 54 and leucine 56 selected as target residues, using the default design parameters. For the MotifGrafting approach, the removed terminal helix was used as the motif that was grafted onto recipient scaffolds using the Rosetta MotifGraft protocol^44^. The scaffold database was comprised of small proteins (<90 amino acids) containing globular proteins from the Protein Data Bank (PDB)^45^ and experimentally determined folds^46,47,48,49^. Residues corresponding to 96, 99, 100, 103 of PIPS2 were selected as hotspot residues on the motif, with residue 96 being an alanine. Following MotifGrafting, a Rosetta Fastdesign protocol was performed to optimize the interface^50^.

Designs were filtered based on computed binding energy (ddG) (≤-50), SASA (≥1300 Å²), shape complementarity (≥0.70), number of interface H-bonds (≥5), and RMSD ≤0.5 Å. The 1685 designs that passed the filter were run through ProteinMPNN (soluble weights) to generate 25 sequences per design. The interface was also re-designed to increase interface diversity. The top 5 sequences determined by ProteinMPNN global score were run through AF2 multimer via ColabFold (v1.5.2)^40^ in single sequence mode with three recycles. Designs were filtered by having a pLDDT > 80, and ipTM > 0.7. 2000 designed binders were ordered as an oligopool from Twist Bioscience, including 783 BindCraft designs and 1217 designs from the MotifGrafting approach.

### Fluorescence activated cell sorting of phosphoPIPS2 Binders

The amplified oligo pool was transformed together with the pCTcon2 vector containing a C-terminal HA tag. For sort 1, induced yeast were washed and labelled with 200nM of either PIPS2 or phosphoPIPS2, containing a His-tag, for 1 hour at 4°C. Cells were washed and labelled with mouse anti-his (MA121315, Invitrogen; 1:100 dilution) for 30 min at 4°C. Cells were washed and labelled with FITC-conjugated goat anti-HA tag antibody (as described above) and PE-conjugated goat anti-mouse antibody (as described above) for 30 min at 4°C. PhosphoPIPS2 samples were sorted based on PE signal. For the second sort, gates were set for both binding and non-binding for both PIPS2 and phosphoPIPS2 50nM samples. Each gate collected 10,000 yeast.

### Library sequencing

After sorting, yeast were cultured in SDCAA media and plasmids were extracted using the Zymoprep Yeast Plasmid Miniprep II (Zymo Research) following the manufacturer’s instructions. The genes of interest were amplified out by PCR, using vector specific primers. A second PCR was performed to add on Illumina adapters and Nextera barcodes. PCR products were purified using the Qiaquick PCR purification kit (Qiagen). Next generation sequencing was performed on an Illumina MiSeq system with 500 cycles, yielding between ∼1.0 million reads per sample. Sequences were translated to the correct open reading frame and matched to their protein sequences. Sequences with a minimum of 5,000 reads that showed a 10-fold enrichment in the phosphoPIPS2 binding gate compared to both the phosphorylated non-binding gate and the non-phosphorylated binding gate were ordered to be tested as single clones.

### Single clone phosphoPIPS2 binder validation

Each design was cloned into a pCTcon2 plasmid (with a C-terminal HA-tag). Induced yeast were washed, and treated with the indicated concentrations of PIPS2 or phosphoPIPS2 for 1 hour at 4°C. Cells were washed and labelled with mouse anti-his (as described above) for 30 min at 4°C. Cells were washed and labelled with FITC-conjugated goat anti-HA tag antibody (as described above) and PE-conjugated goat anti-mouse antibody (as described above) for 30 min at 4°C. Cells were washed, and resuspended in appropriate volumes of 0.1% BSA/PBS before being analyzed.

### Protein expression and purification

Each gene, encoding a C-terminal 6xHis-tag, was purchased from Twist Bioscience, cloned into the pET11 bacterial vector by Gibson assembly and transformed into HB101 *E. coli*. Plasmids were purified using a GeneJET plasmid Miniprep kit and sequences were verified by Sanger sequencing (Microsynth). For bacterial expression, verified plasmids were transformed into T7 Express E. coli. A single colony was picked and grown in LB/Amp as a pre-culture overnight at 37°C. The next day, pre-cultures were used to inoculate expression cultures 1:50, AIM (Auto Induction Media) was used to express all proteins. Cultures were grown at 37°C until the OD600 reached about ∼0.60, at which point they were transferred to 18°C and grown overnight.

The next morning cultures were centrifuged at 4000xg for 15 minutes and the supernatant was decanted. Pellets were resuspended in lysis buffer (50 mM Tris, pH 7.5, 500 mM NaCl, 5% glycerol, 1 mg ml−1 lysozyme, 1 mM PMSF and 1 µg ml−1 DNase) and lysed by sonication. Cell lysates were centrifuged at 20,000xg for 20 minutes and the supernatant was filtered through a 0.22µm filter (99722; Techno Plastic Products). Clarified lysates were loaded onto an ÄKTA pure system (GE healthcare) Ni-NTA HisTrap affinity column followed by size exclusion chromatography on a Superdex HiLoad 16/600 75pg column using PBS. Proteins were concentrated using 3kDa Amicon filters (UFC800396; Sigma), aliquoted, flash frozen with liquid nitrogen and stored at –80°C.

For PKA, the murine catalytic subunit α gene was ordered (sequence in Supplementary Table 1). The expression and purification are identical to the above method, except that TB (Terrific Broth) media was used and the culture was induced with 500µM IPTG before being transferred to 18°C. The PKA protein was concentrated to 2mg/ml in 10kDa Amicon filters (UFC801096; Sigma), then diluted with an equal volume of storage buffer (40mM Tris-HCL, 100mM NaCl, 4mM DTT, 2mM EDTA, 96% glycerol) for a final concentration of 1mg/ml in 48% glycerol and stored at –20°C.

### Preparation of phosphorylated PIPS proteins

For both the phosphoPIPS2 NMR sample and PIPS1-3 phosphorylated samples for CD, MALS, MALDI-TOF and SPR, proteins were incubated in PKA buffer (containing 1mM DTT) with 10-fold molar excess ATP, and a 1000-fold lower molar concentration PKA, relative to PIPS concentrations for 2 hours at 30°C. PIPS4 was phosphorylated as above and buffer exchanged to PBS by serial dilutions using a 3kDa Amicon filter before characterization. For the phosphoPIPS2 NMR sample, the protein was buffer exchanged to 50mM sodium phosphate (pH 6.5) by serial dilutions in a pre-washed 3kDa Amicon filter. For all other samples, proteins were loaded onto a Superdex HiLoad 16/600 75pg column using PBS, then concentrated using 3 kDa Amicon filters.

### Circular Dichroism

Far-ultraviolet circular dichroism spectroscopy measurements were performed on a Chirascan spectrometer (Applied Photophysics). Protein samples were diluted in PBS to a protein concentration of 20µM and transferred to a 1 mm path-length cuvette. Wavelengths between 200 nm and 250 nm were recorded with a scanning speed of 20 nm min−1 and a response time of 0.125 s. All spectra were corrected for buffer absorption. Temperature ramping melts were performed from 20 to 90 °C with an increment of 2 °C min−1. Thermal denaturation curves were plotted by the change of ellipticity at the global minimum of each spectrum. The melting temperatures (Tm) were determined by fitting the data to a sigmoid curve equation on GraphPad Prism (10.2.2).

### Size-exclusion chromatography combined with multi-angle light scattering

Size-exclusion chromatography (controlled by Chromeleon software; Thermo Fisher Scientific, v.7.2.10) combined to a multiangle light scattering device (SEC-MALS, miniDAWN TREOS, Wyatt) was performed to determine the oligomeric state and molecular weight of the purified designs in solution. The final concentration of each sample was approximately 1 mg/ml in PBS (pH 7.4), and 100 μl of each sample was injected into a Superdex 75 10/300 GL column (GE Healthcare) with a flow rate of 0.5 ml/min. UV280, refractive index (dRI) and light scattering signals were recorded. Molecular weight was determined using the ASTRA software (version 6.1, Wyatt).

### Surface plasmon resonance

Affinity measurements were performed on a Biacore 8K (GE Healthcare, software v4.0.8.19879) using HBS-EP+ as a running buffer (10 mM HEPES at pH 7.4, 150 mM NaCl, 3 mM EDTA, 0.005% v/v Surfactant P20, GE Healthcare). PIPS2 and phosphoPIPS2 were immobilized on a CM5 chip (GE Healthcare #29104988) via amine coupling to reach ∼500 response units (RU). PIPS2-B1 and B2 were then injected in serial dilutions using the running buffer. A flow rate of 30 μL/min for a contact time of 120 s was followed by 600 s of dissociation time. Steady-state response units (RUs) were normalized by min-max normalization. Kds were determined by nonlinear four-parameter curve fitting using GraphPad Prism (10.2.2).

### Mass spectrometry

Non-phosphorylated and phosphorylated PIPS samples were diluted 10-fold in MilliQ water and mixed 1:1 (v/v) with sinapinic acid in 0.1% TFA. Samples were spotted onto a 384-well ground steel target plate and measured on a MALDI-TOF AutoFlex Speed mass spectrometer (Bruker) operated in linear positive mode, calibrated for a mass range of 1-20 kDa. Raw data was normalized by min-max normalization and plotted using GraphPad Prism (10.2.2).

### NMR Spectroscopy

All NMR experiments were carried out in a Bruker spectrometer operating at 800.13 MHz ^1^H frequency and equipped with a CPTC ^1^H,^13^C,^15^N 5 mm cryoprobe controlled via an Avance Neo console. All experiments on PIPS and its variants were carried out with a minimum of 300 µM protein solutions prepared in 50 mM sodium phosphate buffer at pH6.5, at 298 K. All spectra were acquired and processed with Bruker TopSpin 4 using the standard pulse sequences as listed, and analyzed with CARA, NMRtist/Cyana, Sparky-NMRFAM and custom scripts as detailed below.

### Preparation of protein samples for solution NMR

A plasmid encoding PIPS2 was transformed into T7 *E. coli* and grown overnight on LB agar plates containing 100µg/ml ampicillin. The next morning, a single colony was cultured in 3mL LB containing ampicillin and grown during the day shaking at 37°C. That evening 1mL of culture was sub-cultured into 30mL of M9 media containing ^15^N (NLM-467-5; Cambridge Isotope Laboratories) and ^13^C (CLM-1396-5; Cambridge Isotope Laboratories) and grown overnight. The next morning the culture was sub-cultured into 1L M9 media containing ^15^N and ^13^C, grown at 37°C shaking until the optical density at 600nm (OD600) hit 0.70, then induced with 1mM IPTG and grown overnight at 18°C. The next day the protein was purified as previously described. Protein samples for NMR spectroscopy were prepared in 50 mM sodium phosphate buffer pH 6.5 with 10% ^2^H_2_O.

### Protein resonance assignment and structure determination

Experiments for backbone resonance assignment consisted in standard triple resonance spectra^51^ HNCA, HN(CO)CA, HNCO, HN(CO)CA, CBCA(CO)NH and HNCACB acquired on a 900 µM sample prepared from cultures grown in M9 medium supplemented with ^13^C-enriched glucose and ^15^N-enriched ammonium chloride. HCCH-TOCSY and ^13^C-resolved NOESY experiments were recorded on the same sample used for backbone assignments; they served for side chain assignment and structure determination together with HNHA, ^15^N-resolved NOESY and ^15^N-resolved TOCSY spectra acquired on a ^15^N-labeled sample plus 2D NOESY and 2D TOCSY spectra collected on an unlabeled sample. Spectra for backbone assignments were acquired with 40 increments in the ^15^N dimension and 128 increments in the ^13^C dimension and processed with 128 and 256 points using forward linear prediction. The HCCH-TOCSY was recorded with 128 increments in the 13C dimensions and processed with 256 and 512 points. ^15^N-resolved NOESY and TOCSY spectra were acquired with 40 increments in ^15^N dimension and 128 in the indirect ^1^H dimension and processed with 128 and 256 points respectively. ^1^H-^1^H 2D-NOESY and 2D TOCSY spectra were acquired with 512 increments in the indirect dimension, processed with 2048 points. Mixing times for NOESY spectra were 120 or 200 ms, and TOCSY spin lock times were set at 60 ms.

All spectra were acquired and processed with Bruker’s TopSpin 4.0 using the standard pulse programs with sensitivity improvement. For PIPS2, backbone assignments were obtained in parallel through manual analysis with the program CARA^52^ and automatically with NMRtist^53^, a webserver implementation of the ARTINA^54^ pipeline that uses a deep neural network to identify signals in NMR spectra, FLYA^55^ for resonance assignment, and if required also automatic NOE identification and analysis for structure determination using CYANA (see original NMRtist papers^56^ and example hands-on applications to protein resonance assignment and structure calculation^57,58,59^). Both procedures returned the same ^1^H,^15^N assignments for backbone units, which covers 85% of the sequence leaving out the flexible termini (residues 1, 2, C-terminal his-tag, and two prolines). Given NMRtist’s accuracy on backbone assignment on our spectra, fully matching our manual assignment, we then ran it on the full set of spectra to obtain automated side chain assignments and structure determination. Resonance assignments were obtained for 92% of all active nuclei, 83% (76% of the full length) being of high confidence. Missing assignments and low-confidence assignments map mainly to the two N-terminal and six C-terminal residues plus Lys41, His42, Tyr43, Lys60 and the stretch Thr85-Arg94; however, the number of assignments available for this region was sufficient to solve the structure even around this region, albeit with a slightly increased local uncertainty, as explained next. NMRtist’s structure calculation procedure converged in the program’s two independent runs to the same structure, indicative of sufficient data quality even at the region with partial assignments. We took Proposal 1 as the final structure, an ensemble calculated from 2047 nOe restraints (503 intraresidual, 569 sequential, 380 medium-range, and 595 long-range) and 198 torsion angle restraints, with close to 1Å RMSD over residues 1 to 113 relative to the mean structure. More structure determination statistics are provided in Supplementary Table 4.2.

For phosphoPIPS2, reliable backbone assignments were obtained only through manual inspection of the 3D spectra, with some additionally missed assignments over those missing for PIPS2, as explained under results. For the S96A and S96E mutants of PIPS2, backbone ^1^H,^15^N assignments were transferred from those of the non-phosphorylated and phosphorylated states, respectively, leaving out those that were ambiguous.

### ^15^N relaxation and solvent exchange

^15^N T_1_ and T_2_ were measured with 128 ^15^N increments processed with 256 points and a total recycle time of 10 seconds, with the following delays: 8, 64 (twice), 136, 232, 336, 472, 664 (twice), 800, 1200 and 1600 ms (for T_1_) and 16, 32 (twice), 64, 96, 128, 160, 192 (twice), 224, 256 and 288 ms (for T_2_). Spectra were acquired and processed in Bruker TopSpin and then analyzed with the relaxation module in Sparky-NMRFAM.

^15^N CEST experiments were obtained with a B_1_ irradiation field of 13 Hz applied at 0.1-1.5 ppm intervals from 100 to 136 ppm during 400 ms, with 8 scans for a total acquisition time of 3-4 days after which the sample stability was checked by ^15^N HSQC. To analyze the CEST data we first manually screened for crosspeaks showing a response by inspecting the pseudo3D spectrum inside CARA, where it was loaded as a ^1^H,^15^N TOCSY to inspect the irradiation axis on the z axis. The ^1^H and ^15^N resonances of residues showing CEST response were loaded onto a custom Matlab script that uses the rbnmr library to read the intensities along the response profile over the irradiation axis and then fits each profile to a two-state Bloch-McConnell model for chemical exchange.

^1^H/^2^H exchange experiments were carried out by 2 consecutive 10-fold dilution/concentration cycles into 50 mM sodium phosphate buffer prepared in 100% ^2^H_2_O (pHmeter reading 6.1 which corresponds to pD=6.5), using 10 kDa amicon devices. The data reported corresponds to the ratio of crosspeak intensities before and after exchange, each normalized to the intensity of isolated methyl groups in the ^1^H spectrum in order to correct for any differences in concentrations.

### Cell culture, transfection and induction

Human Embryonic Kidney (HEK293T; Invitrogen; Ref: R70007) cells were cultured in Dulbecco’s Modified Eagle Medium (41966-029, Gibco) supplemented with 10% (v/v) FBS (A5256701, Gibco) and 1%(v/v) antibiotic penicillin/streptomycin (15140-122, Gibco). Cells were authenticated by the provider (STR genotyping) and tested negative for mycoplasma contamination (qPCR). Cells were maintained at 37°C with 5% CO2 and passaged every two to three days when cells reached about 80% confluency. Cells were seeded into the inner 60 wells of clear 96-well cell culture plates (655-180, Greiner Bio-One), at 10,000 cells per well 24 hours prior to transfection. Cells were transfected by layering 50 μL from a mixture of 330 μL DMEM, 825 – 850 ng total DNA, and 4.125 μg PEI (24765-1, Polysciences) on top of the media in each well, enough for each 6 well column with a 10% extra margin, as described previously^60^. Cells were incubated overnight, for a minimum of 12 hours. The next morning, media was exchanged with fresh media including the respective inducing agents.

### Cellular split-transcription factor assay

Cells were transfected with 100ng PIPS2-Gal4 combined with 100ng of either PIPS2-B1-RelA or PIPS2-B2-RelA, in addition to 250ng of a secreted NanoLuciferase reporter plasmid containing five Gal4 upstream activation sites (UAS) and 400ng of filler plasmid (pCTcon2). For transfections testing point-mutants, the original constructs were replaced by their corresponding point-mutant variants under identical conditions. As a positive control, cells were transfected with 200ng of fused Gal4-RelA plasmid along with reporter and filler plasmid as described above. For the CREB reporter control, 250ng of secreted NanoLuciferase reporter plasmid (regulated by a CRE promoter) was combined with 600ng filler plasmid.

Transfected cells were induced with either 25µM forskolin (CAY-11018-5; Cayman Chemical), 10µM H89 (CAY-10010556-5; Cayman Chemical), or the equivalent amount of DMSO as a vehicle control. After 24 hours, 5µl of medium was transferred to a black 384-well plate (3820, Corning). Each well was then mixed with 5µl of diluted substrate from the Nano-Glo Luciferase Assay kit (N1120, Promega). After gentle shaking, luminescence was measured on a Tecan Spark plate reader with an integration time of 1,000 ms. For statistical analysis, data underwent a Shapiro-Wilk test to determine normality. Data was then analyzed using an unpaired two-tailed Welch’s t-test to determine significance.

### Cell-free reporter assay

The gene encoding the 6xHis-PIPS2-T7 RNA polymerase (T7RNAP) was cloned into a pQE30 plasmid. The plasmid was then transformed into NEBExpress Iq competent *E. coli* (NEB, C3037I) for protein expression. Bacteria were precultured overnight and inoculated to a 500 ml Luria-Bertani (LB)-medium culture, grown until the OD600 was approximately 0.7 and then induced with 0.1 mM IPTG for 3 h. The cells were collected by centrifugation at 4,000g and lysed by sonication. Proteins were purified using Ni-NTA IMAC Sepharose gravity columns. The ZF438-binder fusion protein was expressed using a PURExpress kit from NEB (E6800S). The reaction volume was 10 µl, containing 4 µl of solution A, 3 µl of solution B, 2 µl of DNA template (10 ng µl−1) and 1 µl of water. The reaction was incubated at 34 °C for 3 h and used for the following reporter reaction.

A PURExpress kit from NEB (E6800S) was used to set up the Pepper RNA aptamer reporter expression as well. HBC620 was used as the ligand for Pepper aptamer. The reporter-expressing reaction included 100 nM purified PIPS2-T7RNAP and ZF-binder pre-expressed with PURExpress. The DNA template for the Pepper reporter gene was set to 10 nM, and the gene was transcribed under the regulation of a truncated T7 promoter^61^ downstream of the ZF binding site, which requires a ZF for activation of transcription. Then, 10-µl reactions with different conditions were loaded into a 384-well plate. The Pepper fluorescence intensity was measured on a BioTek Synergy H1 Multimode Reader (Agilent) with an excitation wavelength of 565 nm and an emission wavelength of 615 nm at 30°C with 2-min intervals. In the three-stage experiment, the following enzymes were added: 0.5 µl of 56nM PKA in stage 1, 0.5 µl of 133 U/µl LPP in stage 2, and 0.5 µl of 560nM PKA in stage 3. For control conditions without enzyme, 0.5 µl of the corresponding protein buffer (matching the composition of the enzyme solution) was added. For statistical analysis, the fluorescent yield of each sample was calculated as the difference in RFU values between the first and last timepoint. These values underwent a Shapiro-Wilk test to confirm normality, then analyzed using an unpaired two-tailed Welch’s t-test to determine significance.

## Author contributions

S.B. and Y.M. contributed equally to this work. S.B., L.S., S.J.M. and B.E.C. conceived the project. S.B and B.E.C. designed experiments and experimental methodology. S.B. and M.I. performed the experimental work. S.B. and Y.M. performed the computational work and protein design. S.B. and M.I. performed the expression, purification, and characterization of proteins. L.A.A. and R.R. designed and performed the NMR measurements and NMR structure determination. P.L. performed cell-free assays. S.J.M and B.E.C. provided supervision and acquired the necessary funding. S.B., Y.M., L.A.A, and B.E.C. wrote the manuscript with input from all authors.

## Acknowledgements

We thank the PTPSP facility at EPFL, K. Lau for advice on biophysical characterization of proteins, SCITAS at EPFL for support of computational resources, FCCF at EPFL for assistance with flow cytometry, and EPFL ISIC’s Mass Spectrometry facility for advice regarding mass spectrometry experiments.

## Code availability

The code for designing PIPS is available on GitHub (https://github.com/LPDI-EPFL/PIPS-Design).

## Supplementary Information

**Supplementary Figure 1:**
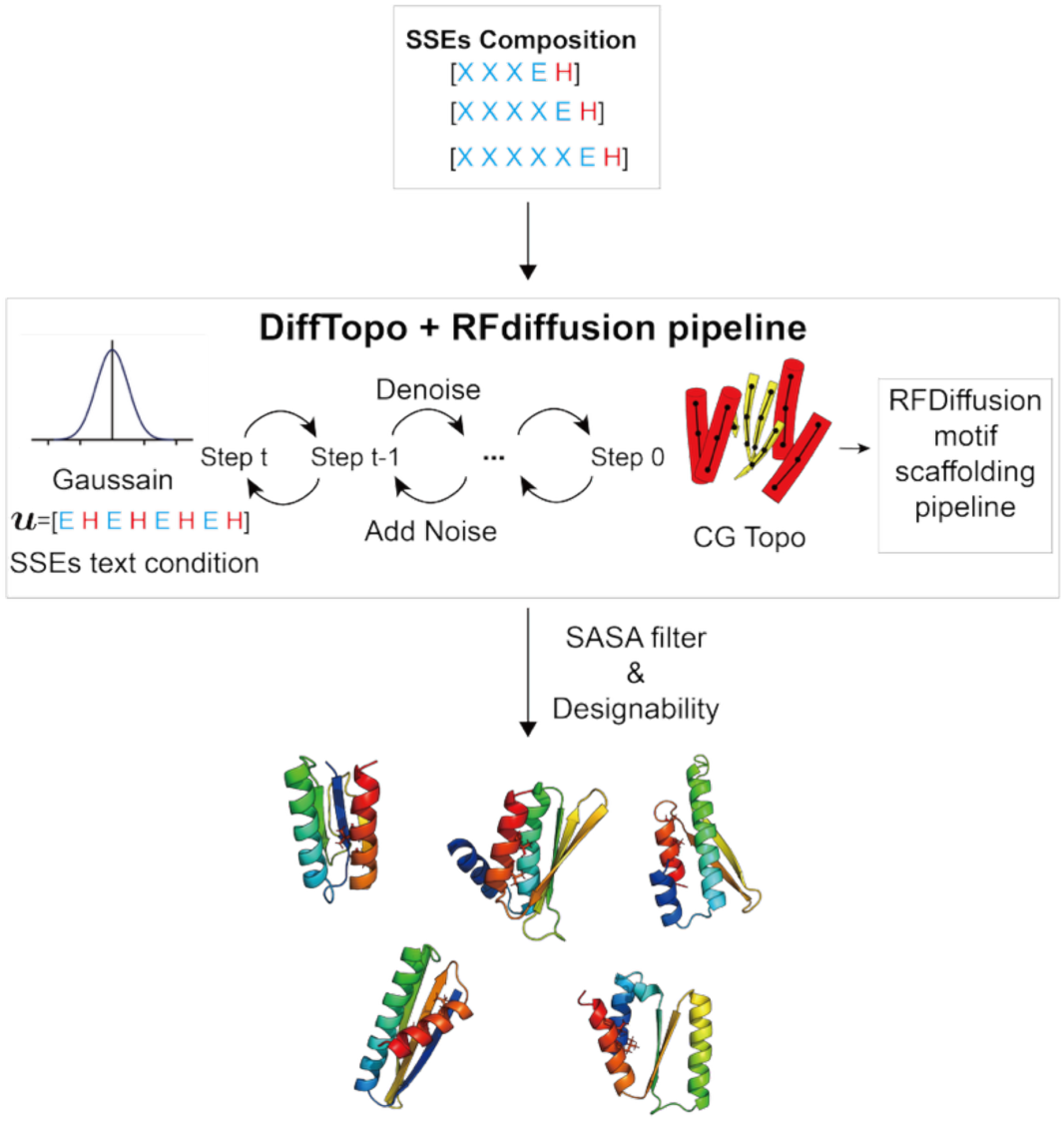
Scaffold diversification and backbone exploration using Difftopo. To identify diverse scaffolds and suitable backbones for transplanting functional sites, we first used a diffusion-based generative model to sample scaffold candidates. Random combinations of secondary structure motifs (with variable elements in the first three positions and a fixed terminal EH motif) were used as input conditions (left block, X ∈ {H, E}, sampled as 3X–5X + EH). These inputs were passed to the topology diffusion model Difftopo to explore feasible fold topologies. The resulting secondary structure element (SSE) patterns were then fed into Difftopo+RFdiffusion, which samples atomic-resolution backbones conditioned on the given SSEs. Finally, we applied SASA-based filtering to identify backbone structures capable of burying a phosphorylation site.

**Supplementary Figure 2.**
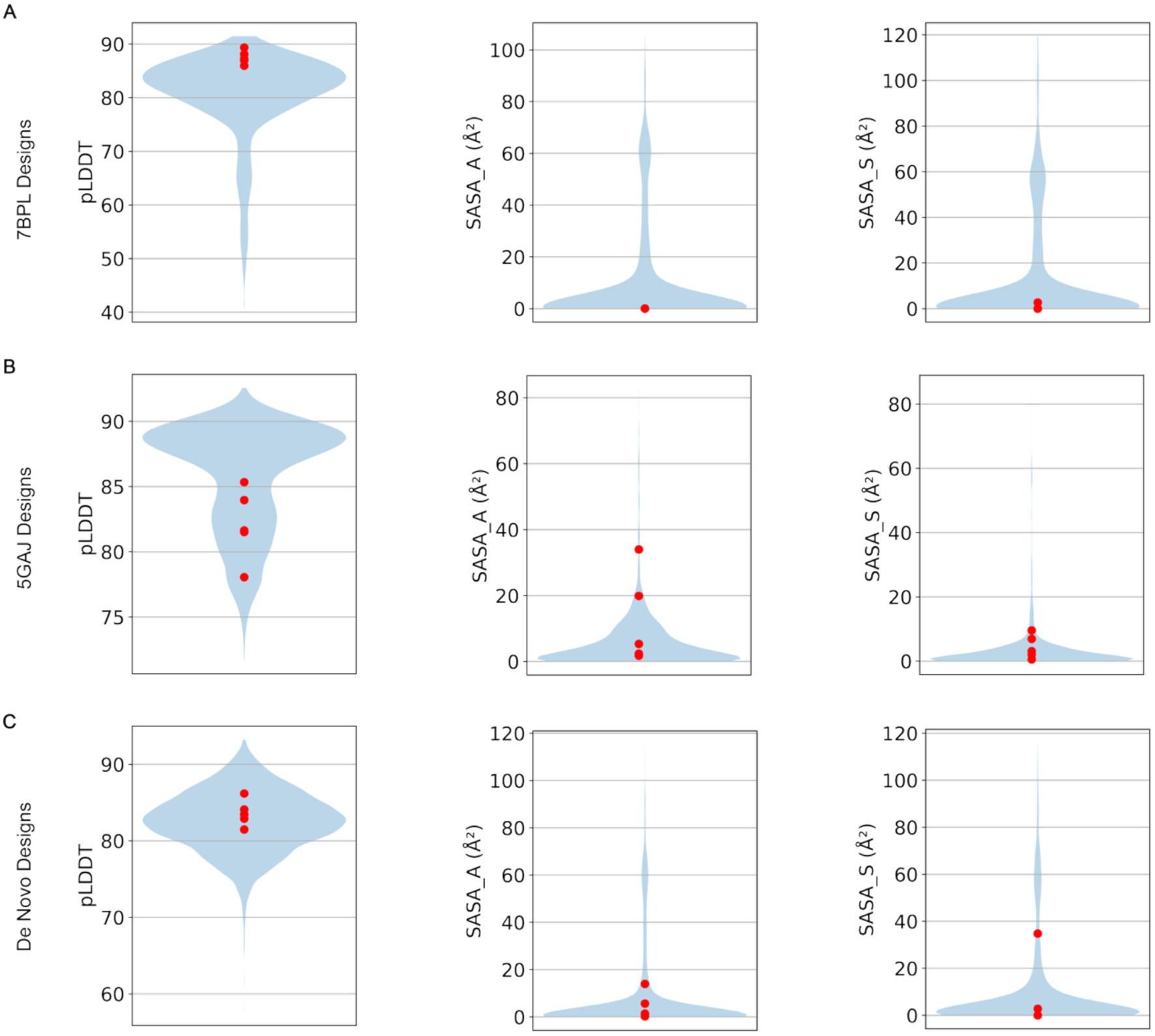
PIPS design metrics. Metrics for PIPS2-4 designs screened in oligopool; including values for pLDDT, SASA of TEV cleavage site alanine, and SASA of the phosphorylation site serine. Values for all 5 AF2 models for each design are included. Red circles represent values for each AF2 model for the validated designs of the **(A)** 7BPL:PIPS2, **(B)** 5GAJ:PIPS3, and **(C)** de novo:PIPS4 scaffolds.

**Supplementary Figure 3:**
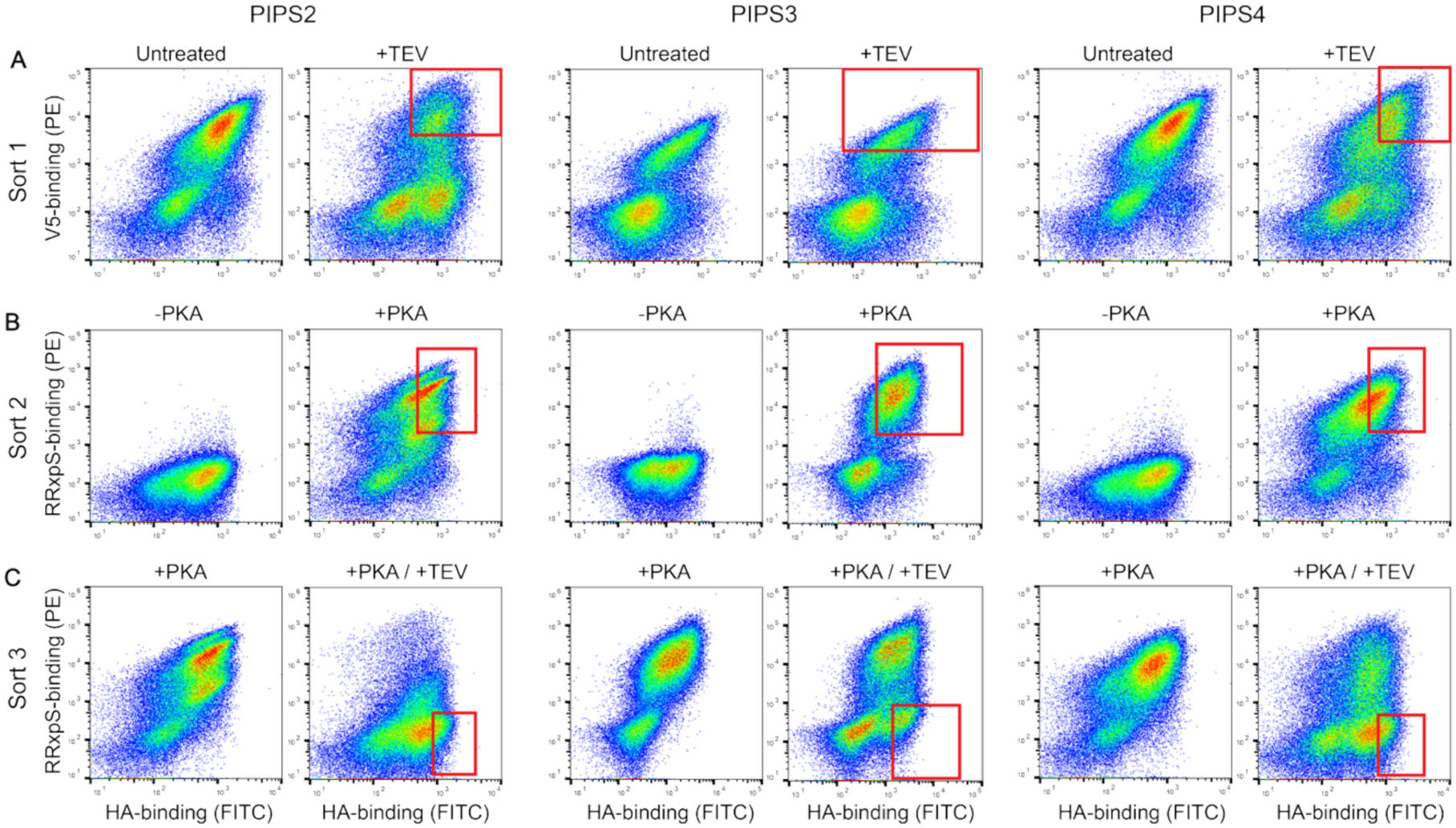
Yeast display screening of PIPS via fluorescence activated cell sorting. Yeast displaying PIPS2-4 were subjected to three rounds of sorting under different treatment conditions. Red rectangles represent gates of sorted yeast. **(A)** Sort 1: Yeast were either untreated or treated with TEV. Yeast were labelled with anti-HA (FITC) to detect surface display (x-axis) and anti-V5 (PE) to detect C-terminal tag retention (y-axis). **(B)** Sort 2: Yest were either untreated or treated with PKA. Yeast were labelled with anti-HA (FITC) (x-axis) and anti-RRxpS (PE) to detect phosphorylation (y-axis). **(C)** Sort 3: Yest were either treated with only PKA or PKA and TEV. Yeast were labelled with anti-HA (FITC) and anti-RRxpS (PE) as in (B).

**Supplementary Figure 4:**
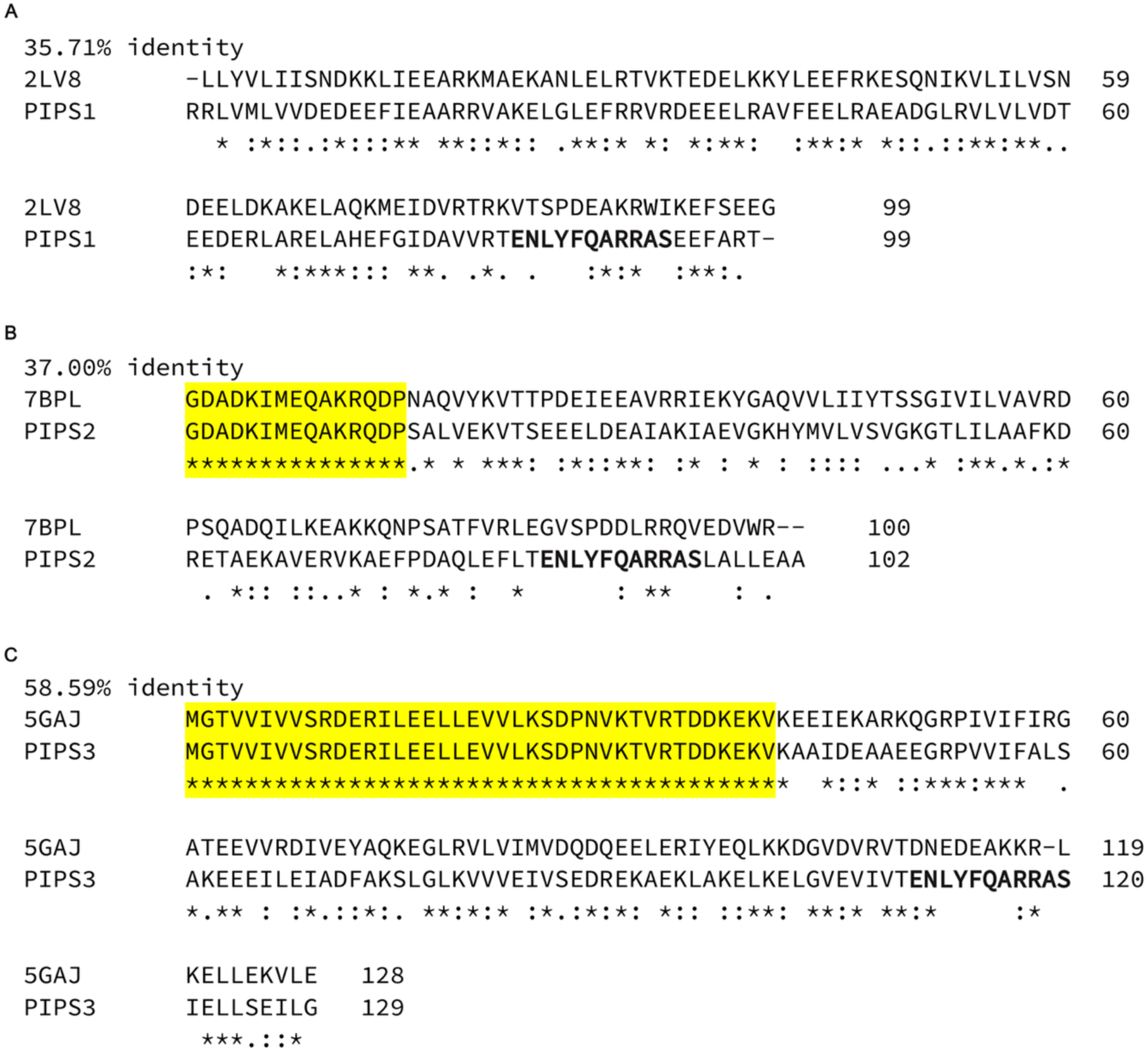
Sequence alignments of validated PIPS1-3 against their initial scaffold. Highlighted regions represent fixed residues during design process. Bold regions represent the phospho-switch motif. Sequence identity percentages are given for each alignment, where the values include the fixed segments. **(A)** PIPS1 aligned with its initial scaffold 2LV8. **(B)** PIPS2 aligned with its initial scaffold 7BPL. **(C)** PIPS3 aligned with its initial scaffold 5GAJ. Alignments were generated using Clustal Omega^62^.

**Supplementary Figure 5:**
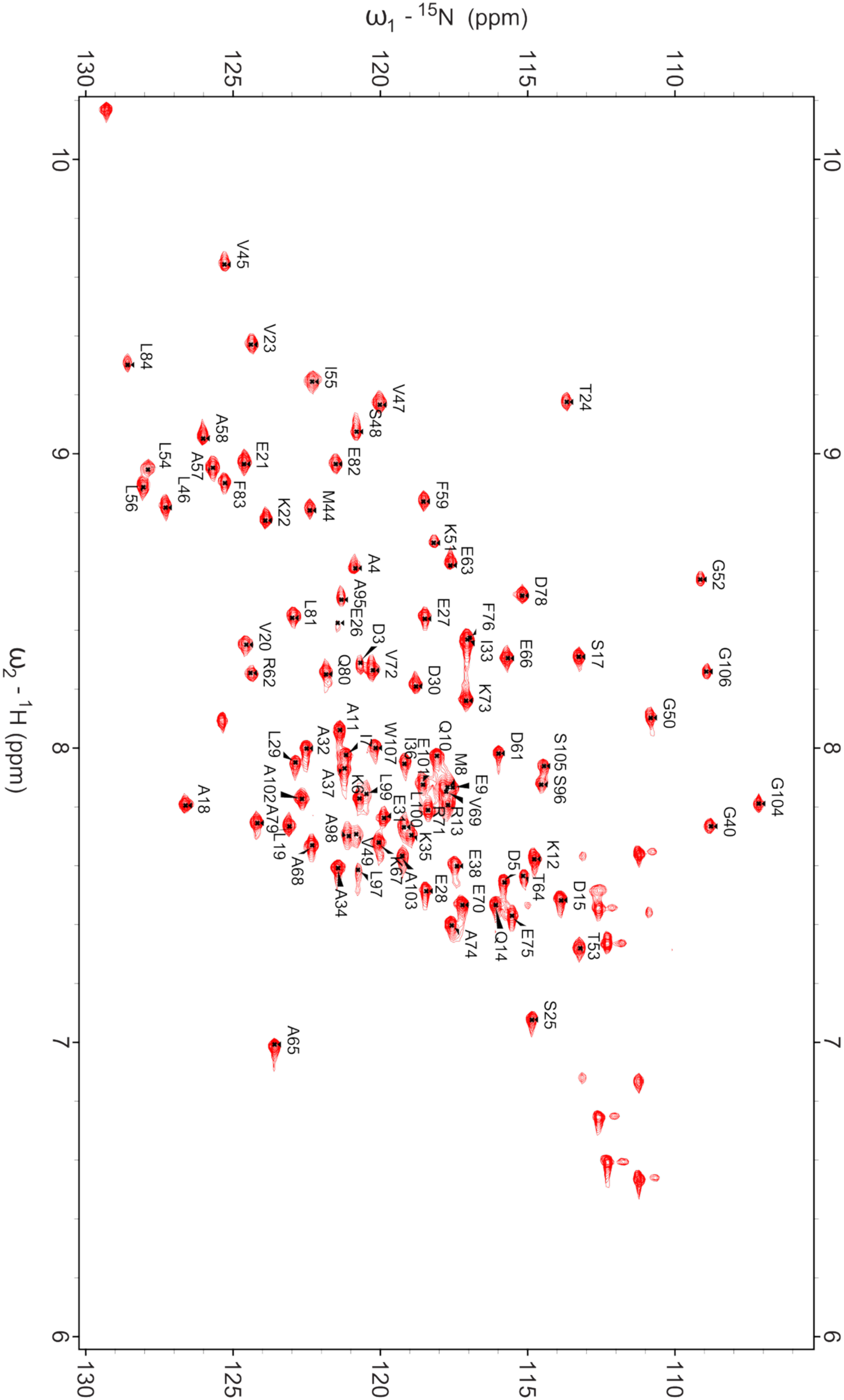
^1^H-^15^N HSQC of non-phosphorylated PIPS2. Backbone assignments are called out of each crosspeak.

**Supplementary Figure 6:**
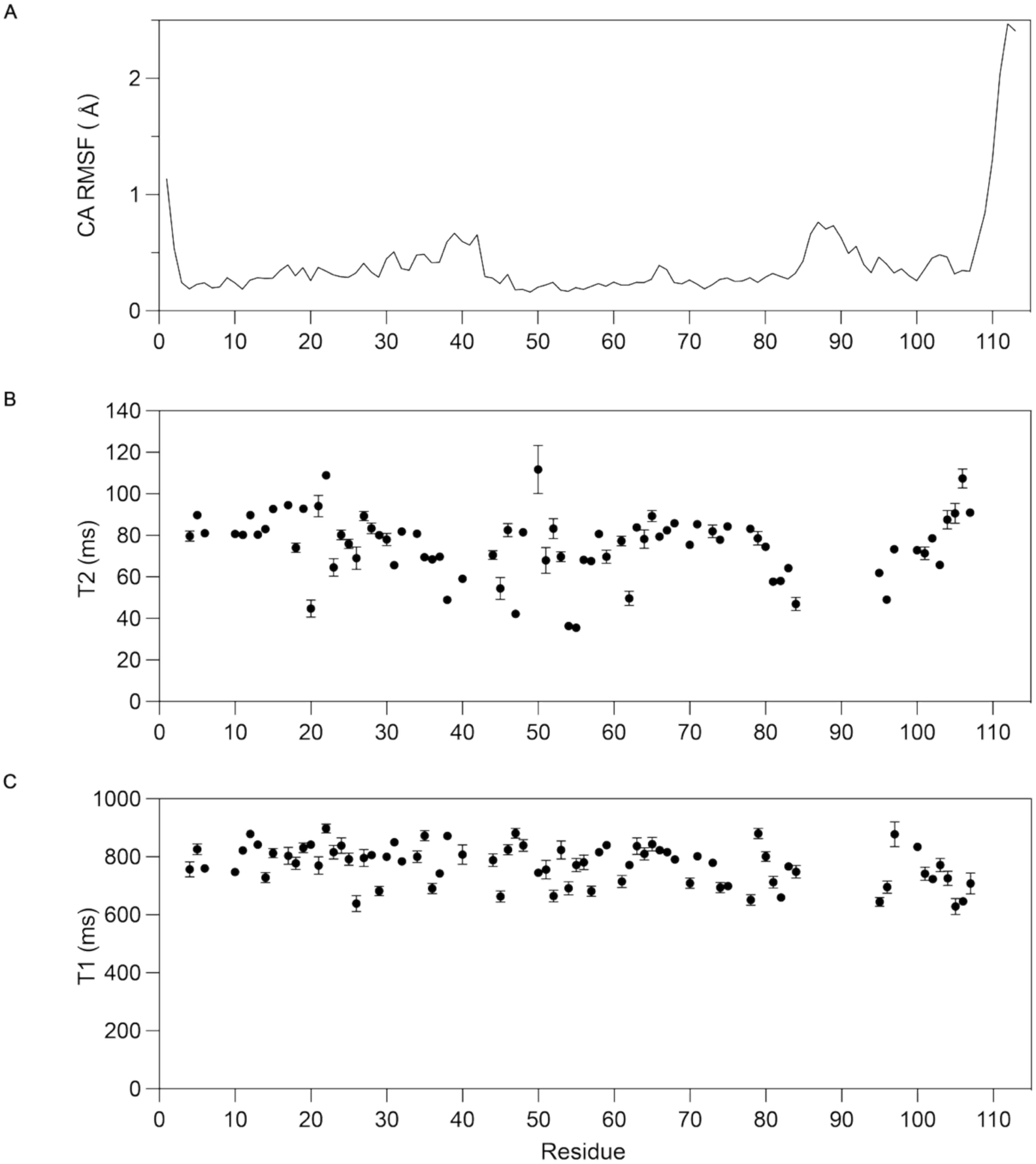
NMR ensemble variability and ^15^N relaxation parameters for PIPS2. **(A)** CA RMSF per residue in the NMR ensemble. **(B)** ^15^N T_2_ relaxation rates. **(C)** ^15^N T_1_. All data at 800.13 MHz ^1^H in phosphate buffer pH 6.5, 298K.

**Supplementary Figure 7:**
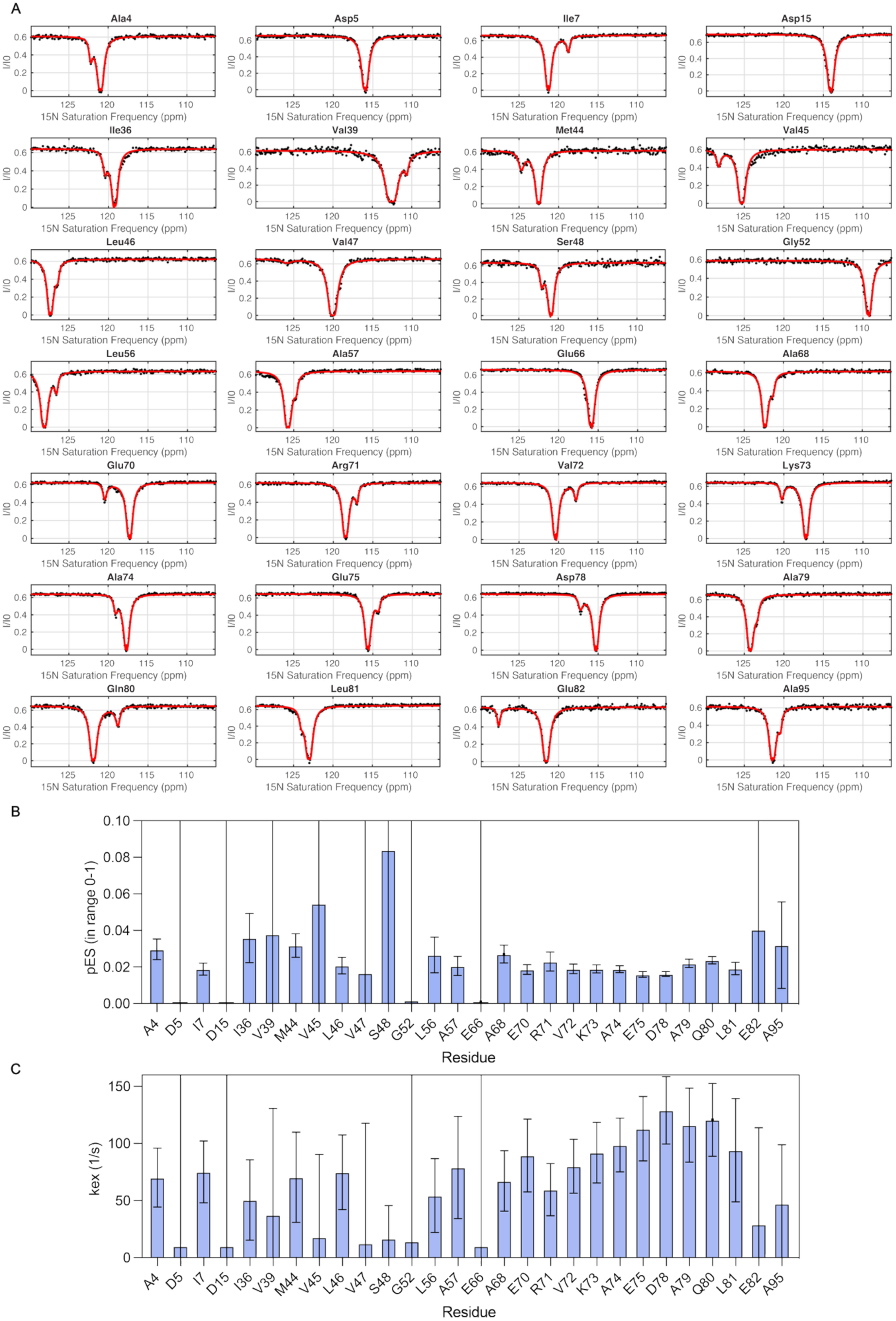
Fits of the ^15^N CEST data for PIPS2 to a two-state model. **(A)** Raw fits on all residues that produced a response for an excited state in the experiment. **(B)** Excited state populations obtained from each residue fit. **(C)** Exchange rates obtained from each residue fit. All data at pH 6.5, 298K, 800.13 MHz ^1^H.

**Supplementary Figure 8:**
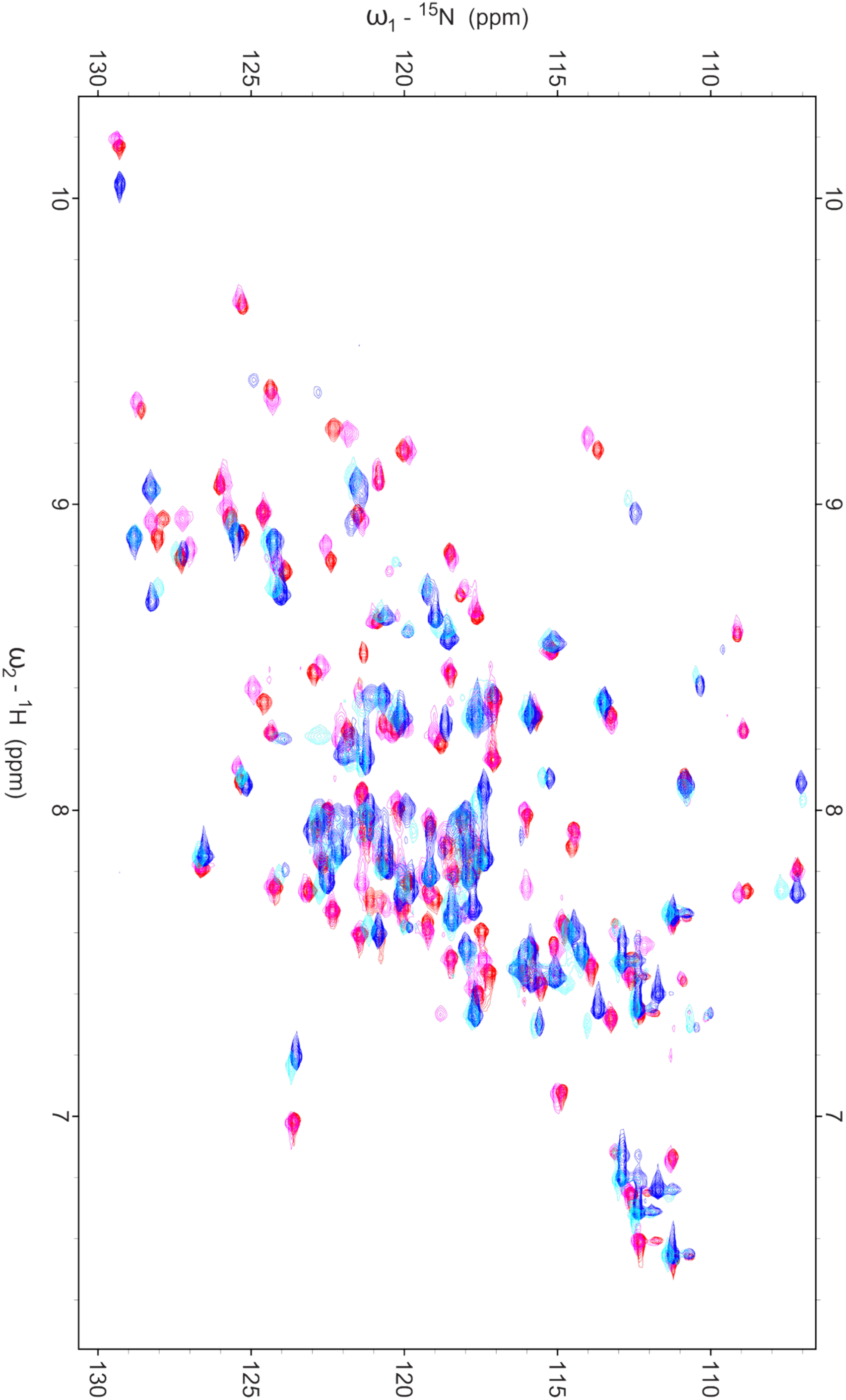
Overlay of ^1^H,^15^N HSQC spectra for PIPS2 variants. Red: PIPS2, pink: S96A mutant, cyan: S96E mutant, and blue: phosphoPIPS2. Data at pH 6.5, 298K, 800.13 MHz 1H.

**Supplementary Figure 9:**
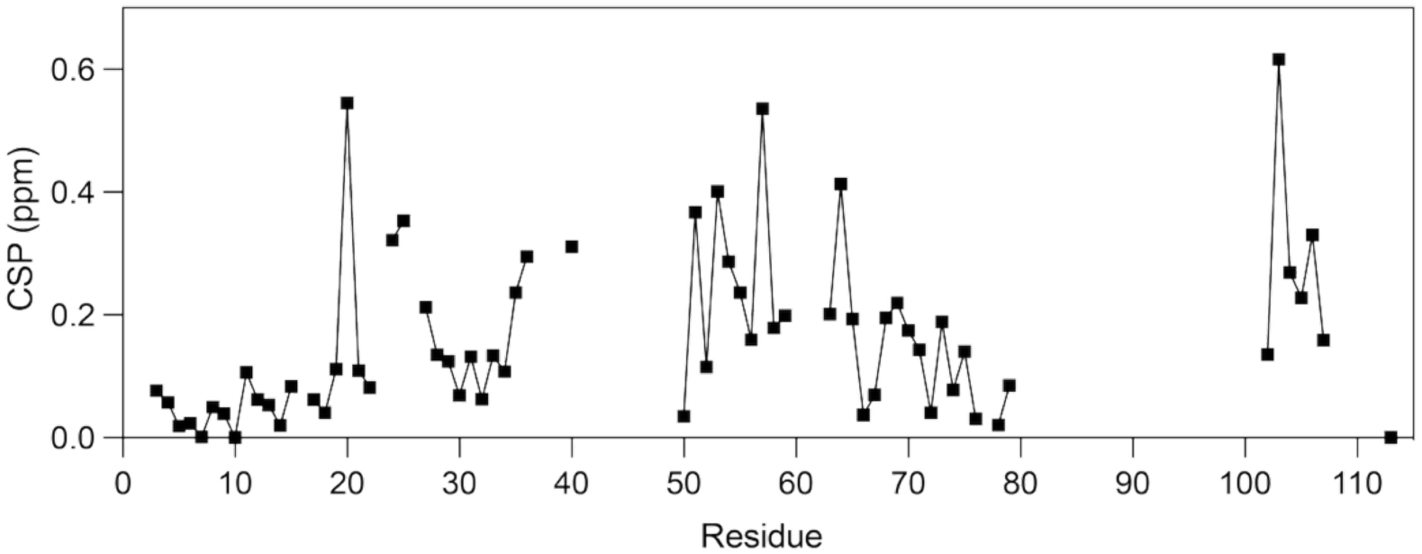
Chemical shift perturbations in phosphoPIPS2 relative to non-phosphorylated PIPS2. Combined ^1^H,^15^N chemical shift perturbation (CSP, in ppm) for residues whose backbone correlations were detected in both the phosphorylated and non-phosphorylated forms. Data at pH 6.5, 298K, 800.13 MHz 1H.

**Supplementary Figure 10:**
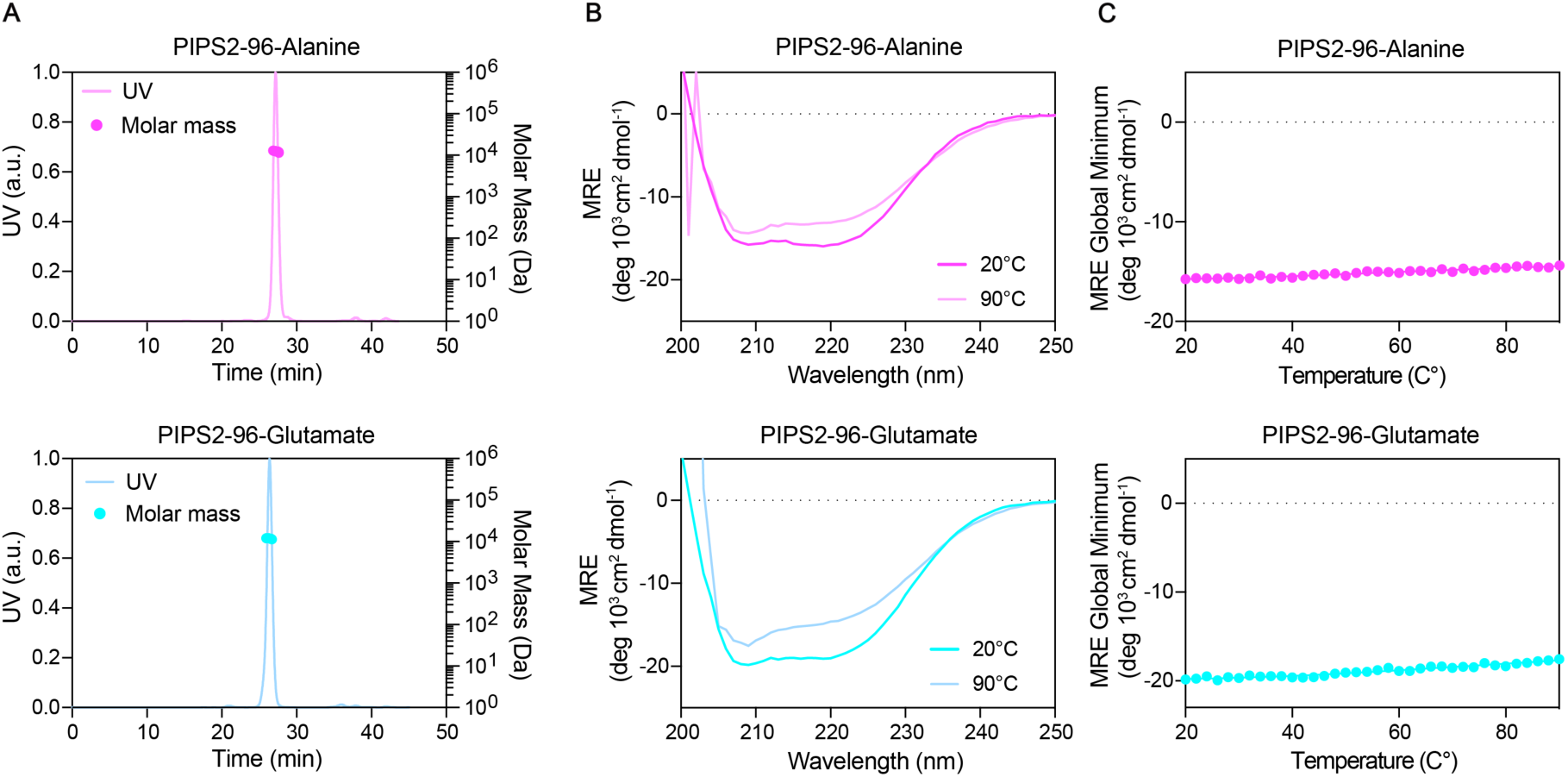
Biophysical characterization of PIPS2 mutants. Variants of PIPS2 containing S96 to alanine and glutamate mutations were characterized. **(A)** Oligomeric state and molecular weight determination by SEC-MALS analysis. **(B)** Protein folding determined by CD at both 20°C and 90°C. **(C)** Thermal stability determined by measuring the MRE global minimum wavelength between 20°C and 90°C.

**Supplementary Figure 11:**
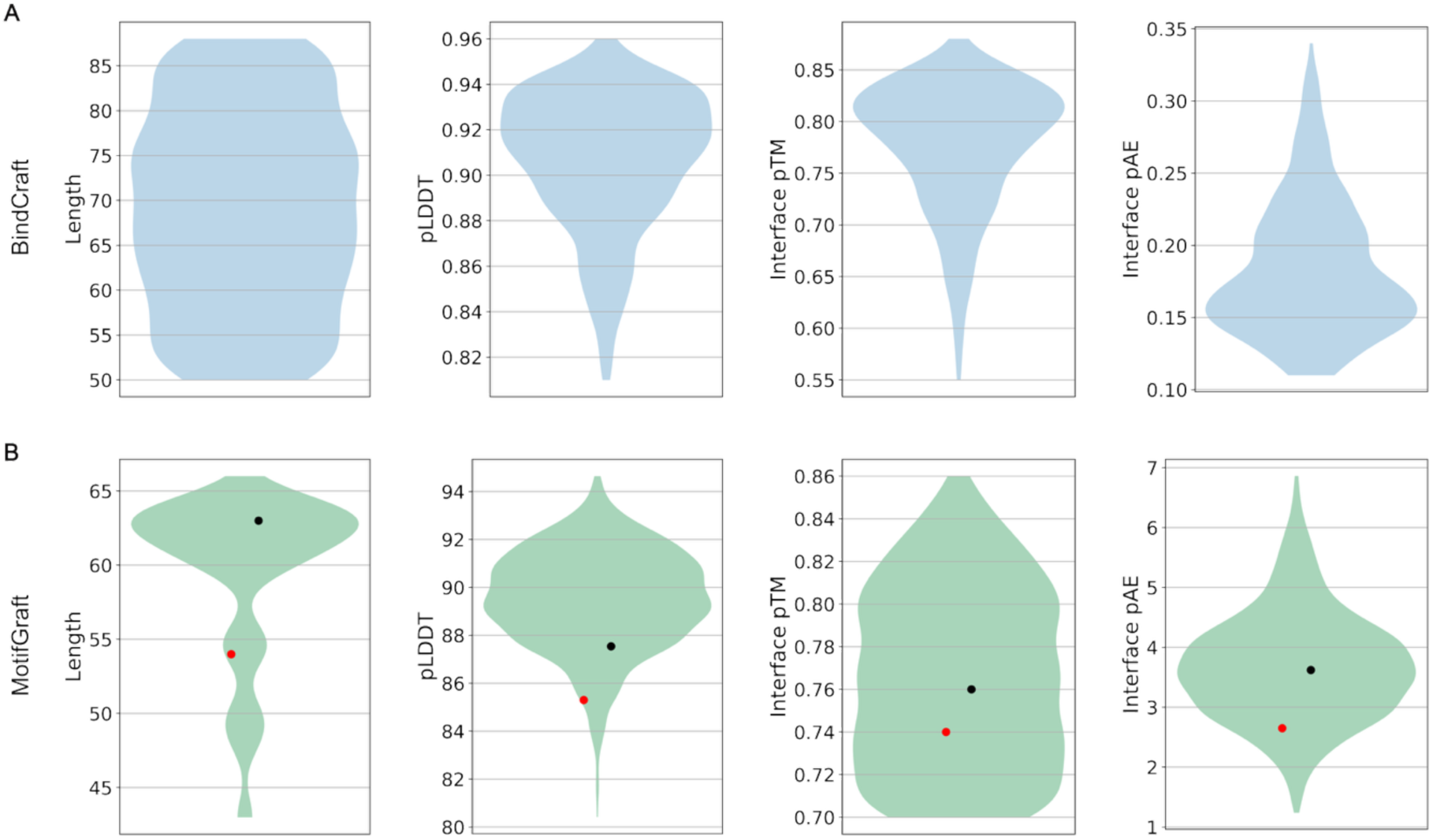
Metrics for designs of phosphoPIPS2 binders screened. The length and AF2-multimer derived binder metrics (pLDDT, ipTM, pTM, and ipAE). Binder metrics were averaged and values plotted for each design. **(A)** BindCraft binder metrics. **(B)** MotifGrafting binder metrics. Red and black circles represent values corresponding to PIPS2-B1 and PIPS2-B2 respectively.

**Supplementary Figure 12:**
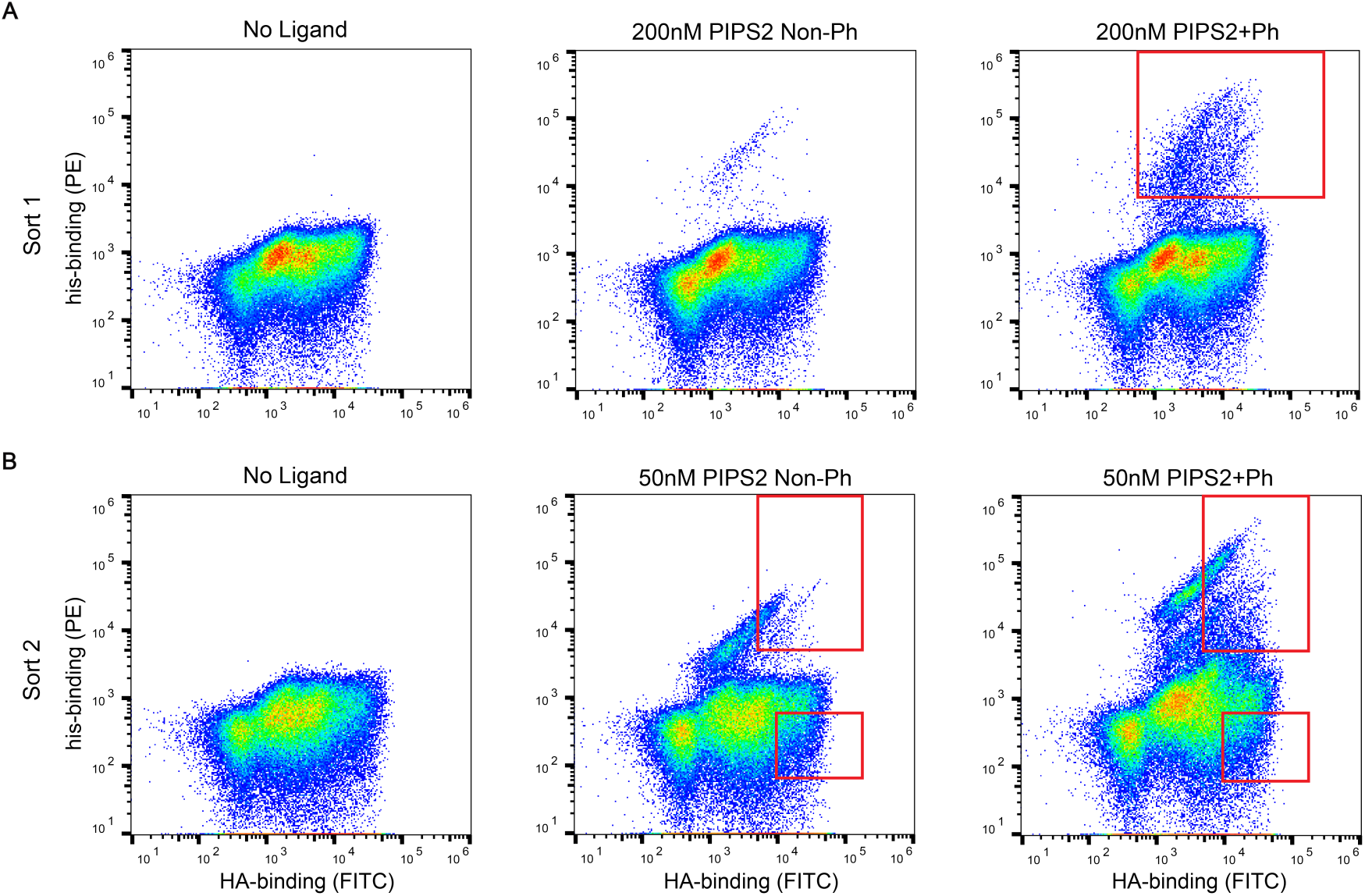
Yeast display screening for phosphoPIPS2 binders via fluorescence activated cell sorting. Flow cytometry data from yeast display screening for phosphoPIPS2 binder designs. Red rectangles represent yeast cells that were sorted. **(A)** Sort 1: Displayed proteins were treated with either no ligand, 200nM PIPS2, or 200nM phosphoPIPS2. Yeast were labelled with anti-HA (FITC) to detect surface display (x-axis) and anti-his (PE) antibodies to detect binding of his-tagged PIPS2 (y-axis). **(B)** Sort 2: Displayed proteins were treated with either no ligand, 50nM PIPS2, or 50nM phosphoPIPS2. Yeast were labelled as in (A). Binding and non-binding populations for both PIPS2 and phosphoPIPS2 were sorted for deep sequencing.

**Supplementary Figure 13:**
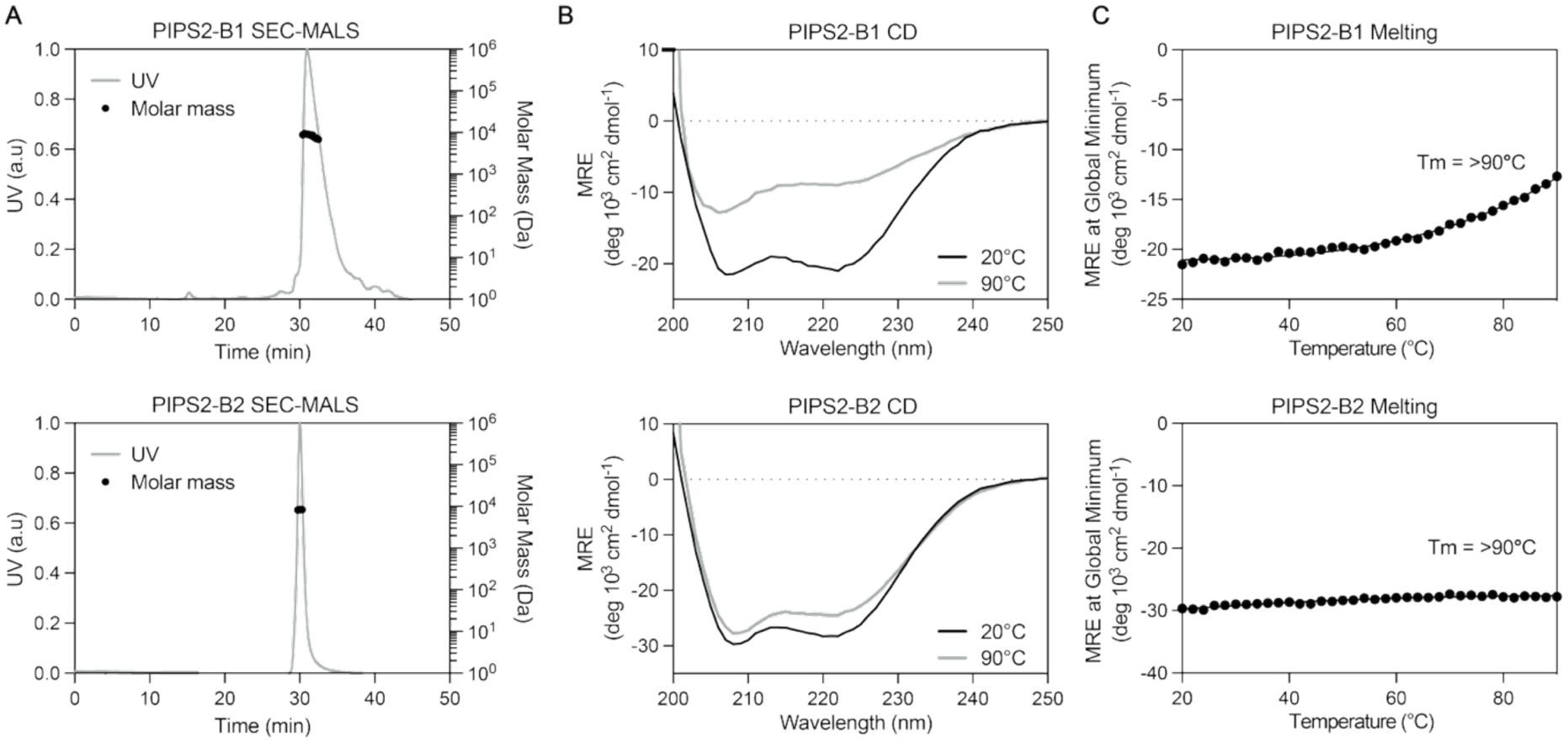
Characterization of purified PIPS2 binders. **(A)** Oligomeric state and molecular weight determination of each binder determined by SEC-MALS. **(B)** CD spectra of each binder measured at 20°C (black) and 90°C (gray). **(C)** Thermal denaturation curves derived from CD spectra, plotted by tracking the global minimum of each spectrum between 20°C and 90°C.

**Supplementary Figure 14:**
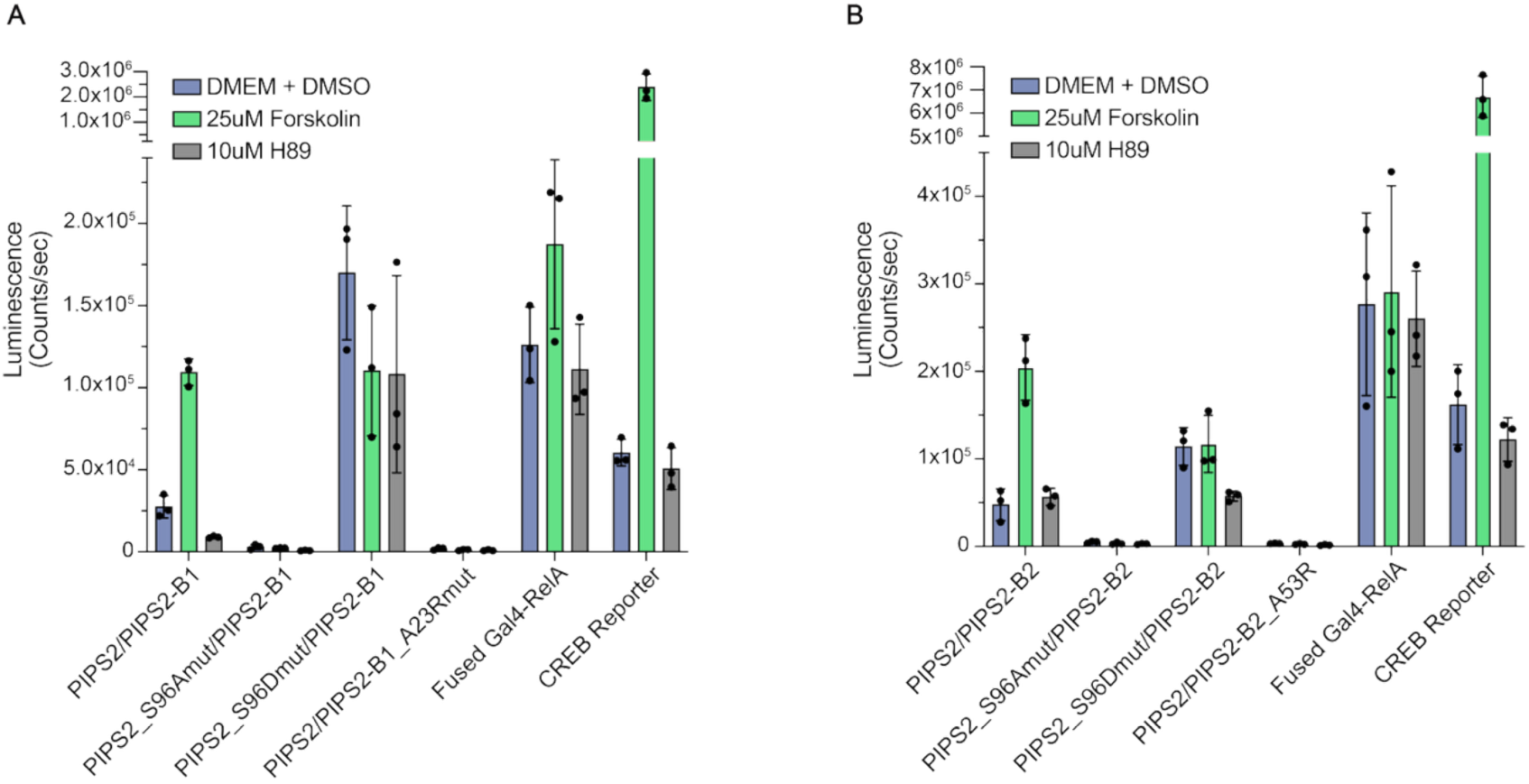
Split transcription factor cell assays. Luminescence measurements for HEK293T cells transfected with PIPS2-Gal4 and either **(A)** PIPS2-B1 or **(B)** PIPS2-B2 fused to RelA. Point mutants of both PIPS2 and binders were used as controls. Cells were either untreated or induced with either 25µM forskolin or 10µM H89. A fused Gal4-RelA construct served as a positive control. A CREB reporter plasmid expressing secreted NanoLuciferase was used as a control to monitor PKA activity.

**Supplementary Figure 15.**
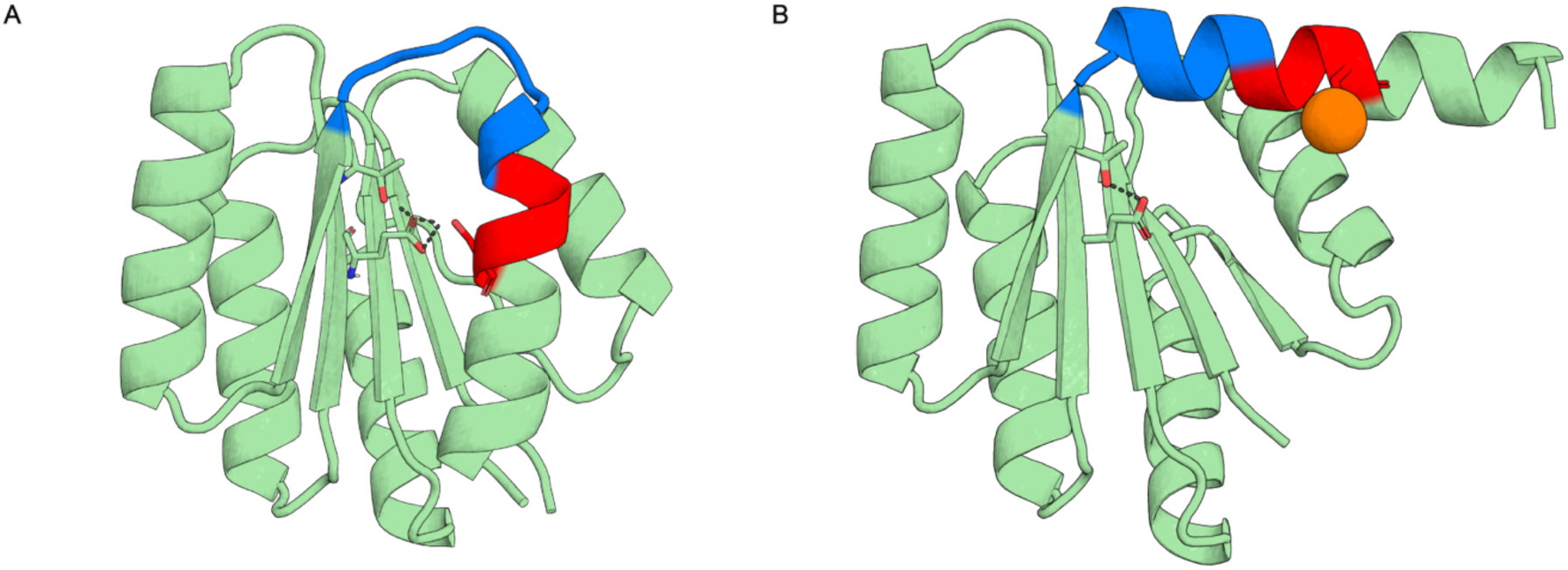
PIPS3 with glutamate in core. **(A)** Original AF2 model of the PIPS3 design. TEV cleavage sequence shown in blue and PKA motif shown in red. Glutamate-83, threonine-109 and serine-120 (phosphorylation site serine) shown as sticks. Dotted black lines represent H-bonds. **(B)** AF3 model of phosphorylated PIPS3, colored as in (A) with phosphate shown as orange sphere.

**Supplementary Figure 16.**
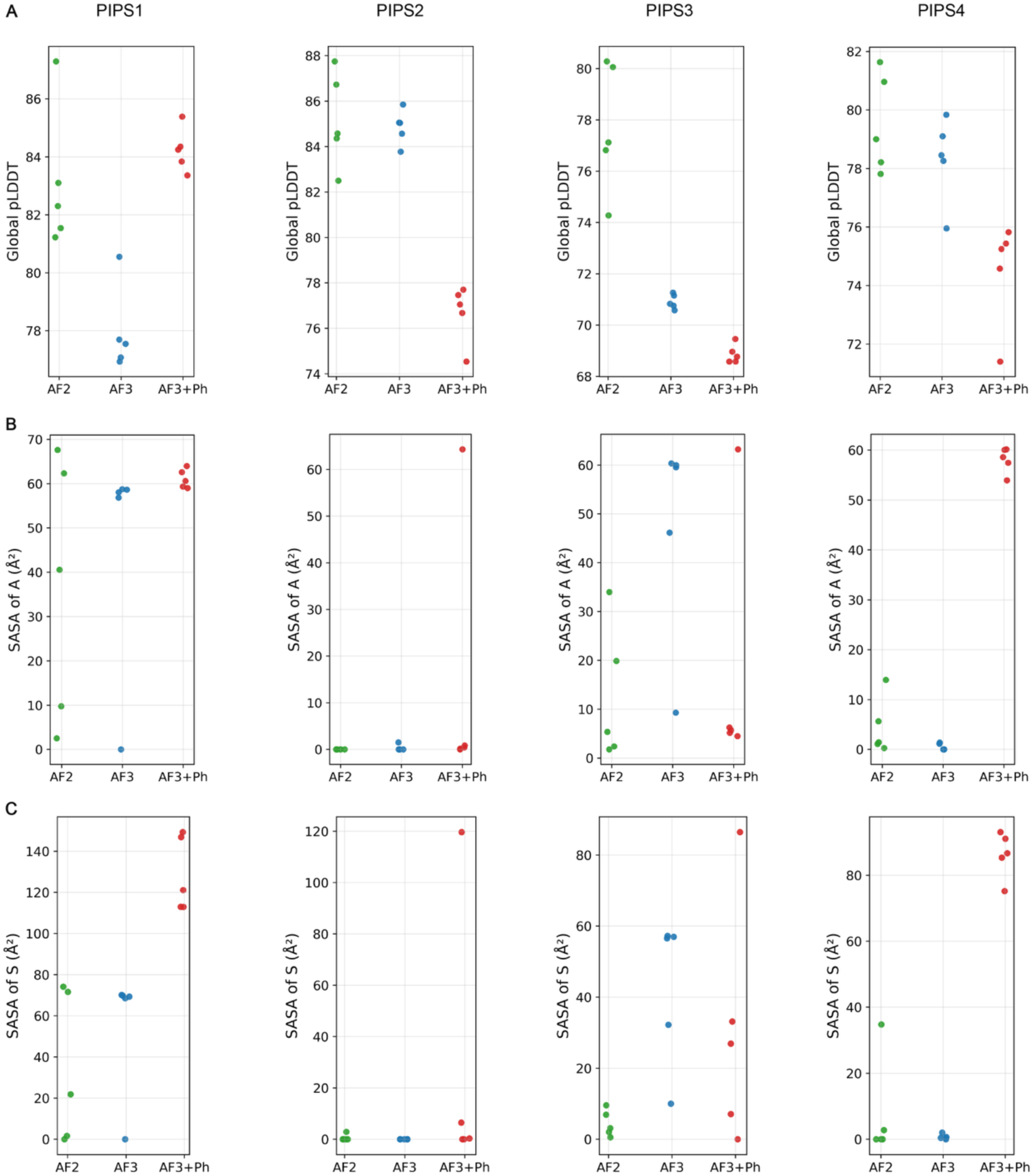
AF2 vs AF3 PIPS metrics. Metrics of all 5 output models of PIPS1-4 from AF2 (green), AF3 (blue), and AF3 with serine of interest phosphorylated (red). **(A)** Global pLDDT values. **(B)** SASA of the TEV cleavage site’s alanine. **(C)** SASA of the phosphorylation site serine.

**Supplementary Figure 17:**
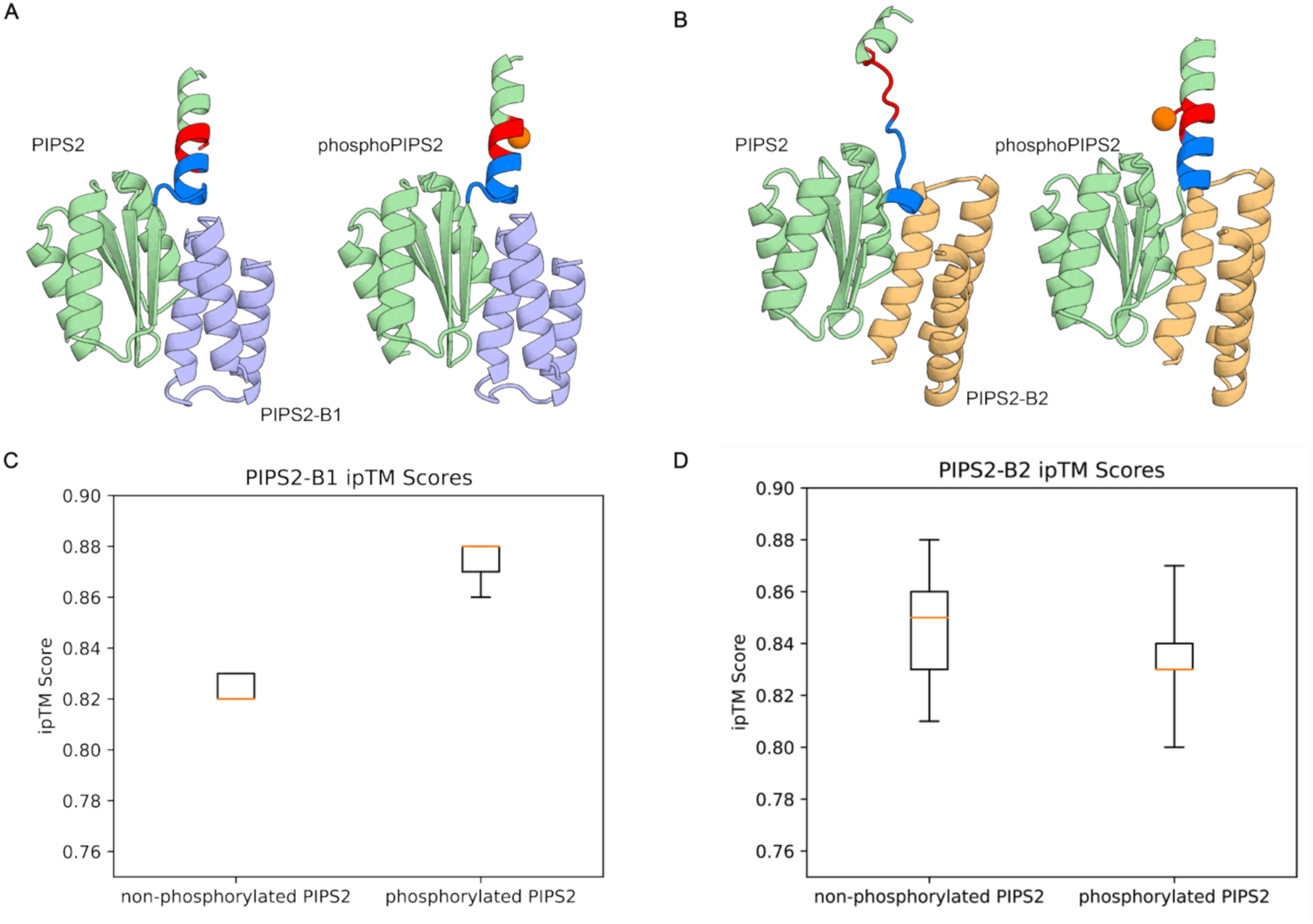
AF3 ipTM comparisons of binders to PIPS2 and phosphoPIPS2. **(A)** AF3-multimer model of non-phosphorylated (pale green, TEV cleavage sequence in blue, and PKA motif in red) and phosphoPIPS2 (phosphate shown as orange sphere) with PIPS2-B1 (light blue). **(B)** AF3-multimer model of PIPS2 and phosphoPIPS2, as colored in (A), with PIPS2-B1 (light orange). **(C)** Comparisons of AF3-multimer ipTM scores for PIPS2-B1 to PIPS2 and phosphoPIPS2. **(D)** Same as (B) but for PIPS2-B2.

**Supplementary Table 1:**
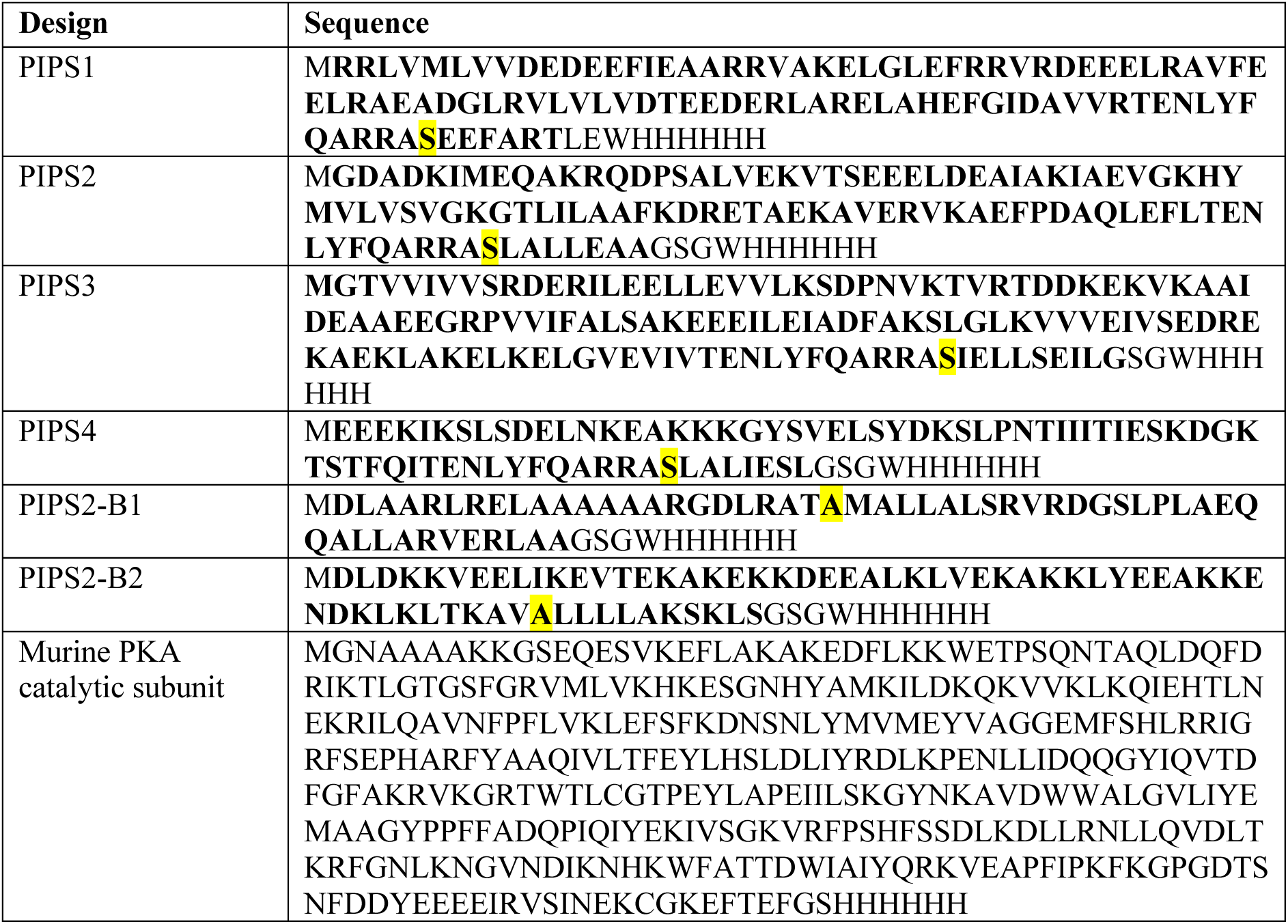
Protein Sequences. Full length sequences of bacterially expressed proteins used for biophysical characterizations. Regions of designed components of each protein in bold. Residues used for mutant variants highlighted.

**Supplementary Table 2:** Refinement statistics of the PIPS2 NMR structure.

|  |  |
| --- | --- |
| **NMR restraints<sup>a</sup>** |  |
| *Total NOEs* | **2047** |
| Intraresidual | **503** |
| Interresidual | **1544** |
| Sequential ($i - j = 1$) | 569 |
| Medium-range ($1 < i - j < 5$) | 380 |
| Long-range ($i - j \geq 5$) | 595 |
| *Torsion angle restraints* | **198** |
| **Structure calculation statistics<sup>a</sup>** |  |
| *CYANA target function value* | 1.93 +/- 0.15 Å<sup>2</sup> |
| *Restraint Violations* |  |
| Distance restraints violations > 0.1 Å | 4 ± 2 |
| Maximal distance restraints violation | 0.17 +/- 0.07 Å |
| Dihedral angle restraint violations > 5° | 0 ± 0 |
| Maximal dihedral angle restraints violation | 3.45 +/- 0.48 ° |
| *Ramachandran plot* |  |
| Residues in most favoured region | 88.3 % |
| Additionally allowed regions | 11.7 % |
| Generously allowed regions | 0 % |
| Disallowed regions | 0 % |
| *RMSD to mean structure through full protein length* |  |
| Backbone<sup>a</sup> | 0.53 +/- 0.14 Å |
| All heavy atoms<sup>a</sup> | 0.99 +/- 0.16 Å |
| *Structure Quality Factors* from Procheck run by PDBsum |  |
| Average Procheck G-factor for dihedral angles | -0.08 |
| Average Procheck G-factor for main chain bond lengths and angles | 0.7 |

<sup>a</sup> From NMRtist run.

## References

1. Borowicz, P., Chan, H., Hauge, A. & Spurkland, A. Adaptor proteins: Flexible and dynamic modulators of immune cell signalling. Scand J Immunol 92, e12951 (2020).

2. Chen, H. et al. Cracking the Molecular Origin of Intrinsic Tyrosine Kinase Activity through Analysis of Pathogenic Gain-of-Function Mutations. Cell Rep 4, 376–384 (2013).

3. Radhakrishnan, I. et al. Solution structure of the KIX domain of CBP bound to the transactivation domain of CREB: a model for activator:coactivator interactions. Cell 91, 741–752 (1997).

4. Somsen, B. A., Cossar, P. J., Arkin, M. R., Brunsveld, L. & Ottmann, C. 14-3-3 Protein-Protein Interactions: From Mechanistic Understanding to Their Small-Molecule Stabilization. ChemBioChem 25, e202400214 (2024).

5. Ardito, F., Giuliani, M., Perrone, D., Troiano, G. & Muzio, L. L. The crucial role of protein phosphorylation in cell signaling and its use as targeted therapy (Review). Int J Mol Med 40, 271–280 (2017).

6. Zhang, Y. et al. 14–3-3ε: a protein with complex physiology function but promising therapeutic potential in cancer. Cell Commun Signal 22, 72 (2024).

7. Pennington, K. L., Chan, T. Y., Torres, M. P. & Andersen, J. L. The dynamic and stress-adaptive signaling hub of 14-3-3: emerging mechanisms of regulation and context-dependent protein–protein interactions. Oncogene 37, 5587–5604 (2018).

8. Aravind, L., Anantharaman, V. & Iyer, L. M. Evolutionary connections between bacterial and eukaryotic signaling systems: a genomic perspective. Current Opinion in Microbiology 6, 490–497 (2003).

9. Nair, A., Chauhan, P., Saha, B. & Kubatzky, K. F. Conceptual Evolution of Cell Signaling. Int J Mol Sci 20, 3292 (2019).

10. Jumper, J. et al. Highly accurate protein structure prediction with AlphaFold. Nature 596, 583–589 (2021).

11. Abramson, J. et al. Accurate structure prediction of biomolecular interactions with AlphaFold 3. Nature 630, 493–500 (2024).

12. Dauparas, J. et al. Robust deep learning based protein sequence design using ProteinMPNN. Science 378, 49–56 (2022).

13. Woodall, N. B. et al. De novo design of tyrosine and serine kinase-driven protein switches. Nat Struct Mol Biol 28, 762–770 (2021).

14. Praetorius, F. et al. Design of stimulus-responsive two-state hinge proteins. Science 381, 754–760 (2023).

15. Guo, A. B. et al. Deep learning-guided design of dynamic proteins. Science 388, eadr7094 (2025).

16. Langan, R. A. et al. De novo design of bioactive protein switches. Nature 572, 205– 210 (2019).

17. Li, P., Martins, I. R. S., Amarasinghe, G. K. & Rosen, M. K. Internal dynamics control activation and activity of the autoinhibited Vav DH domain. Nat Struct Mol Biol 15, 613–618 (2008).

18. Yu, B. et al. Structural and Energetic Mechanisms of Cooperative Autoinhibition and Activation of Vav1. Cell 140, 246–256 (2010).

19. Yang, X. et al. Engineering synthetic phosphorylation signaling networks in human cells. Science 387, 74–81 (2025).

20. Mishra, D. et al. An engineered protein-phosphorylation toggle network with implications for endogenous network discovery. Science 373, eaav0780 (2021).

21. Koga, N. et al. Principles for designing ideal protein structures. Nature 491, 222–227 (2012).

22. Minami, S. et al. Exploration of novel αβ-protein folds through de novo design. Nat Struct Mol Biol 30, 1132–1140 (2023).

23. Koga, N. et al. Role of backbone strain in de novo design of complex α/β protein structures. Nat Commun 12, 3921 (2021).

24. Miao, Y. & Correia, B. DiffTopo: Fold exploration using coarse grained protein topology representations. 2024.02.01.578456 Preprint at 10.1101/2024.02.01.578456 (2024).

25. Watson, J. L. et al. De novo design of protein structure and function with RFdiffusion. Nature 620, 1089–1100 (2023).

26. Dauparas, J. et al. Robust deep learning based protein sequence design using ProteinMPNN. Science 378, 49–56 (2022).

27. Leigh, J. S. Relaxation times in systems with chemical exchange: Some exact solutions. Journal of Magnetic Resonance (1969) 4, 308–311 (1971).

28. Alderson, T. R. & Kay, L. E. Unveiling invisible protein states with NMR spectroscopy. Curr Opin Struct Biol 60, 39–49 (2020).

29. Wayment-Steele, H. K. et al. Learning millisecond protein dynamics from what is missing in NMR spectra. 2025.03.19.642801 Preprint at 10.1101/2025.03.19.642801 (2025).

30. Pacesa, M. et al. One-shot design of functional protein binders with BindCraft. Nature 1–10 (2025) doi:10.1038/s41586-025-09429-6.

31. Balbi, P. E. M. et al. Mapping targetable sites on the human surfaceome for the design of novel binders. 2024.12.16.628626 Preprint at 10.1101/2024.12.16.628626 (2024).

32. England, C. G., Ehlerding, E. B. & Cai, W. NanoLuc: A Small Luciferase Is Brightening Up the Field of Bioluminescence. Bioconjug Chem 27, 1175–1187 (2016).

33. Taylor, S. S. et al. The Tails of Protein Kinase A. Mol Pharmacol 101, 219–225 (2022).

34. Chijiwa, T. et al. Inhibition of forskolin-induced neurite outgrowth and protein phosphorylation by a newly synthesized selective inhibitor of cyclic AMP-dependent protein kinase, N-[2-(p-bromocinnamylamino)ethyl]-5-isoquinolinesulfonamide (H-89), of PC12D pheochromocytoma cells. J Biol Chem 265, 5267–5272 (1990).

35. Lee, P.-W., Mottaghi, S. S., Lugnier, M. G. & Maerkl, S. J. A T7 RNAP regulatory toolbox for cell-free network engineering and biosensing applications. 2025.06.18.660355 Preprint at 10.1101/2025.06.18.660355 (2025).

36. Lang, B., Streit, J. O., Kriwacki, R. W., Christodoulou, J. & Babu, M. M. Protein Dynamics at Different Timescales Unlock Access to Hidden Post-Translational Modification Sites. 2025.06.25.661537 Preprint at 10.1101/2025.06.25.661537 (2025).

37. Tsytlonok, M. et al. Dynamic anticipation by Cdk2/Cyclin A-bound p27 mediates signal integration in cell cycle regulation. Nat Commun 10, 1676 (2019).

38. Scheller, L., Strittmatter, T., Fuchs, D., Bojar, D. & Fussenegger, M. Generalized extracellular molecule sensor platform for programming cellular behavior. Nat Chem Biol 14, 723–729 (2018).

39. Watson, J. L. et al. De novo design of protein structure and function with RFdiffusion. Nature 620, 1089–1100 (2023).

40. Mirdita, M. et al. ColabFold: making protein folding accessible to all. Nat Methods 19, 679–682 (2022).

41. Zhang, Y. & Skolnick, J. TM-align: a protein structure alignment algorithm based on the TM-score. Nucleic Acids Res 33, 2302–2309 (2005).

42. Chao, G. et al. Isolating and engineering human antibodies using yeast surface display. Nat Protoc 1, 755–768 (2006).

43. Lau, K., Bouchri, B. & Pojer, F. Purification of 10xHis-SuperTEV. (2023).

44. Silva, D.-A., Correia, B. E. & Procko, E. Motif-Driven Design of Protein-Protein Interfaces. Methods Mol Biol 1414, 285–304 (2016).

45. Berman, H. M. et al. The Protein Data Bank. Nucleic Acids Res 28, 235–242 (2000).

46. Rocklin, G. J. et al. Global analysis of protein folding using massively parallel design, synthesis, and testing. Science 357, 168–175 (2017).

47. Linsky, T. W. et al. Sampling of structure and sequence space of small protein folds. Nat Commun 13, 7151 (2022).

48. Tobin, A. R. et al. Inhibition of a malaria host-pathogen interaction by a computationally designed inhibitor. Protein Sci 32, e4507 (2023).

49. Bhardwaj, G. et al. Accurate de novo design of hyperstable constrained peptides. Nature 538, 329–335 (2016).

50. Fleishman, S. J. et al. RosettaScripts: a scripting language interface to the Rosetta macromolecular modeling suite. PLoS One 6, e20161 (2011).

51. Sattler, M., Schleucher, J. & Griesinger, C. Heteronuclear multidimensional NMR experiments for the structure determination of proteins in solution employing pulsed field gradients. Progress in Nuclear Magnetic Resonance Spectroscopy 34, 93–158 (1999).

52. Keller, R. *The* Computer Aided Resonance Assignment Tutorial. (Cantina Verlag, 2004).

53. Klukowski, P., Riek, R. & Güntert, P. NMRtist: an online platform for automated biomolecular NMR spectra analysis. Bioinformatics 39, btad066 (2023).

54. Klukowski, P., Riek, R. & Güntert, P. Rapid protein assignments and structures from raw NMR spectra with the deep learning technique ARTINA. Nat Commun 13, 6151 (2022).

55. Schmidt, E. & Güntert, P. A new algorithm for reliable and general NMR resonance assignment. J Am Chem Soc 134, 12817–12829 (2012).

56. Gottstein, D., Kirchner, D. K. & Güntert, P. Simultaneous single-structure and bundle representation of protein NMR structures in torsion angle space. J Biomol NMR 52, 351– 364 (2012).

57. Giudice, J. et al. Requirements for efficient endosomal escape by designed mini-proteins. bioRxiv 2024.04.05.588336 (2024) doi:10.1101/2024.04.05.588336.

58. Yanaka, S. et al. Identification of potential C1-binding sites in the immunoglobulin CL domains. Int Immunol 36, 405–412 (2024).

59. Paul, J. & Deshmukh, M. V. Chemical shift assignment of dsRBD1 and dsRBD2 of Arabidopsis thaliana DRB3, an essential protein involved in RNAi-mediated antiviral defense. Biomol NMR Assign 18, 99–104 (2024).

60. Scheller, L. Synthetic Receptors for Sensing Soluble Molecules with Mammalian Cells. Methods Mol Biol 2312, 15–33 (2021).

61. Hussey, B. J. & McMillen, D. R. Programmable T7-based synthetic transcription factors. Nucleic Acids Res 46, 9842–9854 (2018).

62. Madeira, F. et al. The EMBL-EBI Job Dispatcher sequence analysis tools framework in 2024. Nucleic Acids Res 52, W521–W525 (2024).

